# Identification of nonsense-mediated decay inhibitors that alter the tumor immune landscape

**DOI:** 10.1101/2023.12.28.573594

**Authors:** Ashley L Cook, Surojit Sur, Laura Dobbyn, Evangeline Watson, Joshua D Cohen, Blair Ptak, Bum Seok Lee, Suman Paul, Emily Hsiue, Maria Popoli, Bert Vogelstein, Nickolas Papadopoulos, Chetan Bettegowda, Kathy Gabrielson, Shibin Zhou, Kenneth W Kinzler, Nicolas Wyhs

## Abstract

Despite exciting developments in cancer immunotherapy, its broad application is limited by the paucity of targetable antigens on the tumor cell surface. As an intrinsic cellular pathway, nonsense-mediated decay (NMD) conceals neoantigens through the destruction of the RNA products from genes harboring truncating mutations. We developed and conducted a high throughput screen, based on the ratiometric analysis of transcripts, to identify critical mediators of NMD. This screen implicated disruption of kinase SMG1’s phosphorylation of UPF1 as a potential disruptor of NMD. This led us to design a novel SMG1 inhibitor, KVS0001, that elevates the expression of transcripts and proteins resulting from truncating mutations *in vivo* and *in vitro*. Most importantly, KVS0001 concomitantly increased the presentation of immune-targetable HLA class I-associated peptides from NMD-downregulated proteins on the surface of cancer cells. KVS0001 provides new opportunities for studying NMD and the diseases in which NMD plays a role, including cancer and inherited diseases.

**One Sentence Summary:** Disruption of the nonsense-mediated decay pathway with a newly developed SMG1 inhibitor with *in-vivo* activity increases the expression of T-cell targetable cancer neoantigens resulting from truncating mutations.

## Introduction

Despite success with cancer immunotherapies, approved immunotherapies are not available for the majority of cancer patients and only a minority of treated patients realize a durable response[1-4]. Current studies largely focus on discovering new agents and identifying patients most likely to benefit from existing immunotherapies[5]. While many studies correlate tumor insertion and deletion (indel) mutation load with immunotherapeutic response, not all tumors with high indel mutational loads respond to checkpoint inhibitors[6-13].

The typical adult solid tumor contains a median of 54 coding somatic nucleotide variants, many of which have the potential to create novel neoantigens or Mutation Associated NeoAntigens (MANA)[14-16]. Approximately 5% of mutations are insertions/deletions (indels) or splice site changes that alter the open reading frame (ORF) of the transcript[14-16]. This subset of mutations is of particular interest because they can result in proteins and derived peptides that are foreign to a host’s healthy cells, giving rise to neoantigens[17-19]. In normal cells, nonsense-mediated decay (NMD) plays an important role in messenger RNA (mRNA) quality control, as well as normal gene expression[20-23]. In cancer cells, however, NMD may aid immuno-evasion by eliminating RNA transcripts coming from genes that carry truncating mutants[19]. This prevents translation and presentation of peptides from these proteins on MHC class I complexes, rendering them invisible to immune cells[24]. Previous work has shown knockdown of the NMD pathway with siRNA enhances the anti-tumor immune response[18, 25].

Analysis of cell lines with loss of the UPF1 RNA helicase and ATPase gene (*UPF1*), a key mediator of NMD degradation, also showed increased levels of aberrant transcripts and mutant proteins in the alleles targeted by NMD[26]. This is reminiscent of the pharmacological modulation of splicing, which has also been shown to increase the number of neoantigens present on the cancer cell surface due to similar underlying mechanisms[27]. Despite early studies showing little toxicity with NMD inhibition, there is as of yet no reports of a specific chemical inhibitor of the pathway with good bioavailability[28-32].

In this work we create a cell-based high throughput assay to query the effects of a curated library of small molecules on NMD function. From this screen, we identify a lead compound capable of inhibiting NMD and ascertain its protein target as nonsense mediated mRNA decay associated PI3K related kinase (SMG1), a kinase that is a critical mediator of NMD. As the lead compound produced unacceptable toxicity in animal models, we then designed a specific and bioavailable small molecule inhibitor of SMG1 (KVS0001) that is well tolerated *in vivo*. We then demonstrate that targeted inhibition of SMG1 by KVS0001 leads to the presentation of novel neoantigens identifiable by T cells leading to tumor growth inhibition *in vitro* and *in vivo*.

## Results

### Development of a high throughput assay to identify NMD inhibitors

To develop an assay to find NMD inhibitors, we identified isogenic cell lines with out-of-frame indel mutations, hereinafter referred to as truncating mutations, targeted by NMD activity (Table S1). We previously reported a panel of non-cancerous cell lines in which 19 common tumor suppressor genes were inactivated using the Clustered Regularly Interspaced Short Palindromic Repeats (CRISPR)-Cas9 system[33]. Through evaluation of this panel, we discovered two genes, Stromal Antigen 2 (*STAG2*) and Tumor Protein p53 (*TP53*), which did not express their expected proteins when assessed by western blots (STAG2) or immunohistochemistry (p53) (Figure S1). Because the inactivation of these two genes was the result of frame-shift mutations, we suspected that the absence of the proteins was due to NMD. This suspicion was supported by whole transcriptome RNA-sequencing (Figure S2)[33]. Notably we saw an average decrease of STAG2 RNA transcripts by 20-fold and TP53 by 6-fold relative to their respective wild-type transcripts in the parental cell lines.

We selected two *STAG2* knockout clones (clone 2 and 8) and one *TP53* knockout clone (clone 221) derived from the Retinal Pigmented Epithelial (RPE1) cell line to design a high-throughput screen (HTS). Treatment with the canonical protein synthesis inhibitor, working indirectly to inhibit NMD, emetine demonstrated up to a 60-fold recovery of mutant RNA transcript expression, establishing this panel as appropriate for HTS (Figure S3)[34].

We then designed a Next Generation Sequencing (NGS) assay for NMD (Figure 1A). We mixed the three cell lines in equal proportions and plated the mixture in 96-wells plates followed by treatment with one compound (drug) per well. We determined NMD inhibition efficacy by comparing the wild-type and truncating mutant transcript expression levels in a ratiometric manner. Specifically, the wild-type sequences of the reciprocally knocked out clone (i.e., wild-type *STAG2* sequence from the *TP53* knockout clones, and wild-type *TP53* sequence from the *STAG2* knockout clones) served as internal references, providing a ratiometric assay of mutant to wild-type transcript abundance. This ratiometric assay minimized confounders introduced by nonspecific transcriptional activators or generally toxic agents.

**Figure 1:**
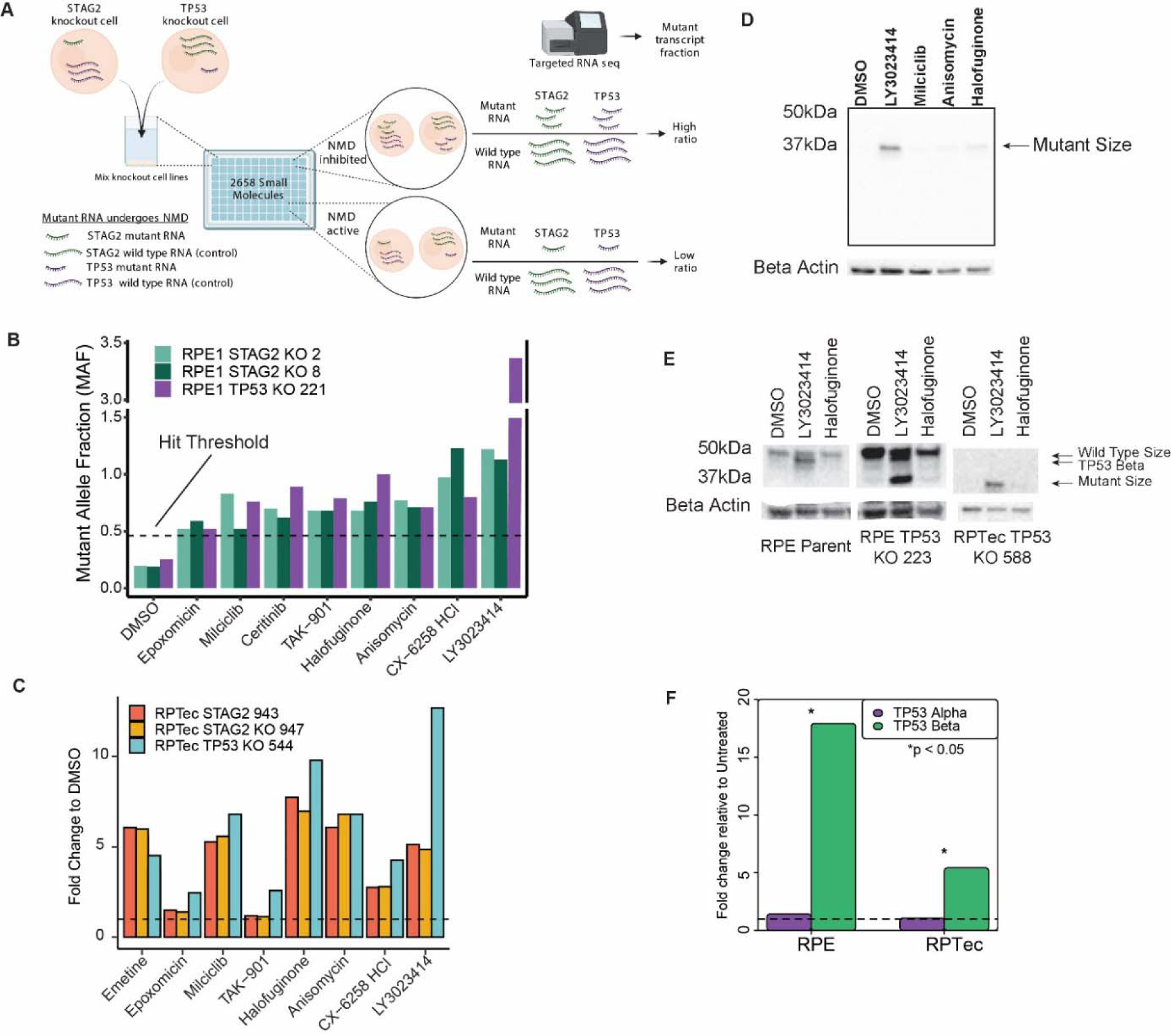
LY3023414 is a small molecule capable of increasing transcription of NMD targets. **A)** Schematic of HTS used to identify inhibitors of NMD. Mutant transcripts are represented by a smaller length in the cartoon for illustrative purposes only. All small molecules were tested at 10μM. **B)** Mutant RNA reads relative to wild-type reads for the top 8 hits from the HTS. The dotted line represents the minimum fraction required to be considered a hit (> 5 standard deviations above DMSO control). Full screen results are presented in Figure S4. **C)** Targeted RNA sequencing results of isogenic RPTec knockout clones treated with the 8 best hits from the HTS at 10µM. The dotted line represents a relative RNA expression level of 1, equal to that of DMSO treated wells. Data for ceritinib, which did not validate on any line, is presented only in Figure S8. **D)** TP53 western blot on RPE TP53 224, containing a homozygous *TP53* mutation, using the four hit compounds that validated in RPTec isogenic lines at 10μM. **E)** Western blot analysis of full-length TP53α and isoform TP53β after treatment with two NMD inhibitor lead candidates at 10μM. TP53β (expression known to be controlled by NMD) as well as mutant TP53 are prominently induced by LY3023414 whereas full-length is not. Note that RPE TP53 223 is a heterozygous knockout clone with one near wild-type allele whereas RPTec TP53 588 contains a homozygous *TP53* indel mutation. **F)** qPCR showing 10μM LY3023414 treatment causes increased expression of the NMD controlled alternative transcript for TP53, TP53β, in parent cell lines for RPE1 and RPTec. Significance determined by Student’s t-test. Unless indicated otherwise cells were exposed to test compound for 16 hours.

The use of three cell lines with different truncating mutations from two different target genes minimized the possibility that drugs identified in the HTS were cell-line clone or mutation-specific. Note that the use of these cell lines, carefully mixed, banked, and preserved, did not substantially increase the amount of time or work required to screen a single cell line. A combination of well and plate barcodes allowed the pooling and scoring of over 1920 assays in a single NGS lane (see methods).

### Execution of a high throughput screen to identify NMD inhibitors

Previous human clinical trials suggest that off-target toxicity at doses required for NMD inhibition makes emetine and other well-known NMD inhibitors unsuitable for human use[35-39]. We performed a high throughput screen (HTS) to identify more specific NMD inhibitors by treating the isogenic cell line panel described above with a commercially available library consisting of 2,658 FDA-approved or in late-phase clinical trial small molecules and natural products (Table S2). After purifying RNA from cells 16 hours post-treatment, we scored NMD inhibition using the strategy described in Figure 1A and Figure S4. Predictably, emetine increased the relative expression of mutant to wild-type transcripts by 3 to 4 fold on average (Figure S5). Eight compounds (0.3% of the library) increased the ratiometric mutant transcript fraction more than five standard deviations above the DMSO controls in all three cell lines (Figure 1B). This hit threshold was chosen as it was the minimum required to ensure no false positives were observed in the DMSO controls. One of these eight compounds, anisomycin, is a known inhibitor of protein synthesis and commonly used NMD inhibitor for *in vitro* studies, providing independent validation of the screen[34]. The other seven compounds increased mutant RNA transcript levels 5 to 10-fold relative to untreated cells.

### LY3023414 inhibits NMD and causes re-expression of mutant RNA and protein

To validate the eight hit compounds described above, we tested their effects in additional lines with mutations targeted by NMD (Table S1). First, we assessed them on isogenic *STAG2* and *TP53* knockouts in RPtec cells, another non-cancerous cell line (Figures S6A, S6B, and S7)[33]. Four of the original eight hit compounds increased mutant RNA expression in a dose-dependent manner (Figures 1C and S8). Next, we examined the effects of these four compounds on additional RPE1 TP53 knockout cell lines with different mutations predicted to generate truncated TP53 proteins (Figures S6C and S7). While treatment with all four potential NMD inhibitors restored expression of the truncated mutant TP53 proteins, two of them (LY3023414 and halofuginone) did so most robustly (Figures 1D and 1E).

Additionally, *TP53* has an isoform, TP53β, whose expression is known to be controlled by the NMD pathway[40]. Using cell lines with intact TP53β isoform transcripts, we observed an increase in the TP53β isoform in cell lines treated with both LY3023414 and halofuginone (Figure 1E middle and left column). Quantitative real time PCR (qPCR) of both the full-length TP53 (TP53α) and NMD sensitive (TP53β) transcripts in parental RPE1 and RPTec cells treated with LY3023414 or halofuginone confirmed the selective upregulation of TP53β, but not TP53α, transcripts (Figure 1F). Based on a consistently stronger effect of LY3023414 over halofuginone across multiple isogenic cell lines and assays, LY3023414 was chosen to be the initial lead compound for further studies involving NMD inhibition.

### LY3023414 increases expression of NMD repressed mutant RNA transcripts and proteins *in vitro* and *in vivo*

To evaluate whether LY3023414 could relieve NMD repression of naturally occurring heterozygous mutant transcripts, we chose the NCI-H358 and LS180 cancer cell lines (Table S1). Treatment with LY3023414 followed by whole transcriptome RNA sequencing revealed increased expression of the mutant allele in 42% and 67% of heterozygous, out-of-frame, indel mutations in these two lines (Figure 2A). A third of the nonsense mutations in NCI-H358 were also “recovered” (i.e., mutant transcripts increased relative to wildtype transcripts) after treatment with LY3023414 (Figure 2A middle). In LS180 there was only one nonsense mutation meeting the required coverage, so it could not be evaluated in depth. Single base pair substitutions not resulting in stop codons were not affected by LY3023414 in either line (Figure 2A right).

**Figure 2:**
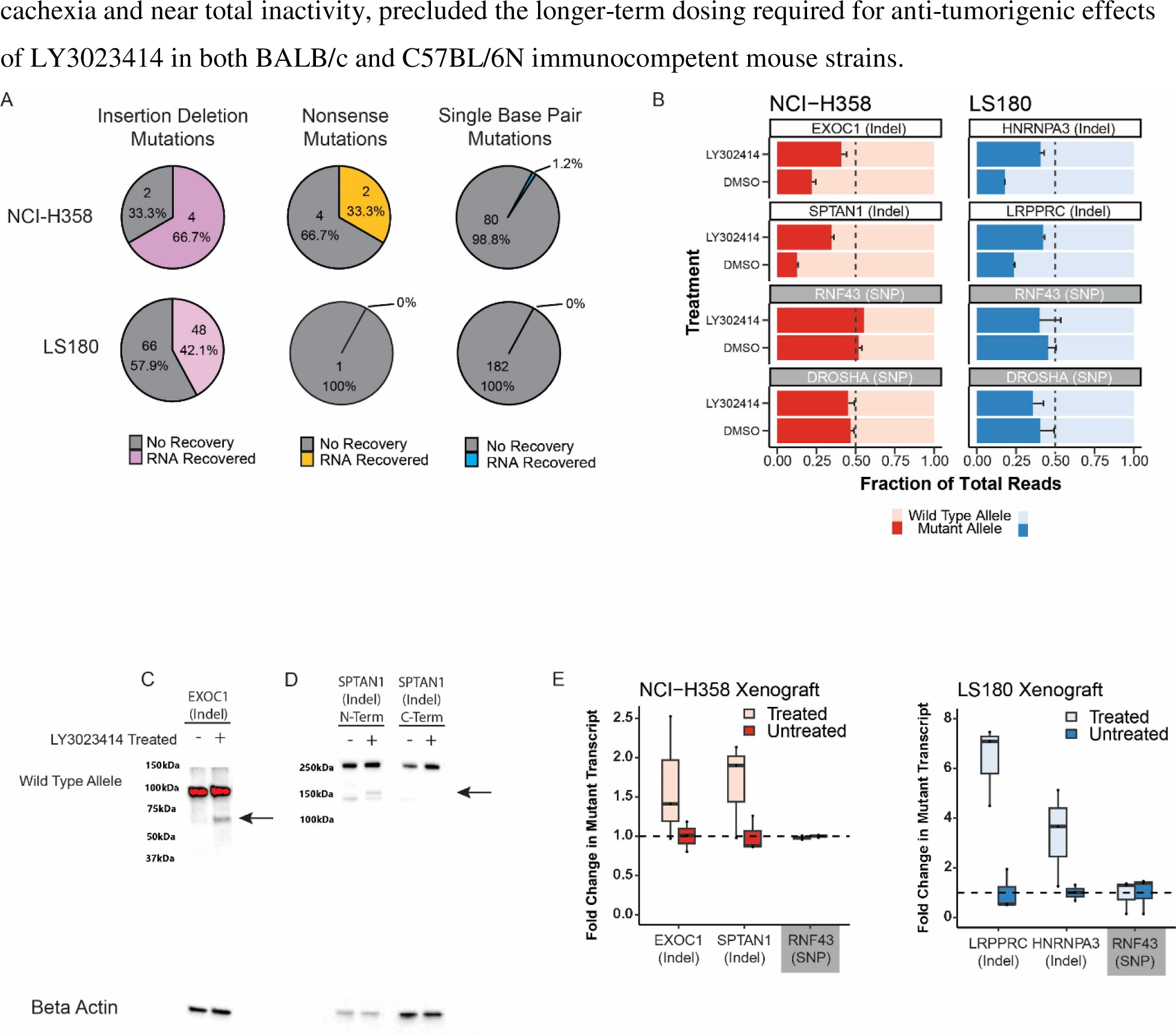
Inhibiting NMD in cancer cells increases broad expression of truncated gene mRNA and protein. **A)** Mutant transcript recovery rates for genes containing heterozygous indel mutations based on RNA-sequencing results in cell lines treated with 5µM LY3023414 for 16 hours. Strict inclusion criteria were used, such that only mutations with sufficient sequencing coverage are shown (see methods). Recovery is defined as at least 2-fold increase over DMSO treatment. **B)** Targeted high coverage RNA-sequencing confirms recovery of mutant transcript levels in NCI-H358 and LS180 cancer cell lines treated with 5µM LY3023414. *RNF43* and *DROSHA* contain common heterozygous SNPs and the mutant allele refers to the non-reference genome allele. Error bars indicate 95% confidence limits. **C)** Western blot analyses of NCI-H358 cells showing mutant and wild-type protein levels in EXOC1 and **D)** SPTAN1 with and without 5µM LY3023414 treatment. The black arrow indicates the expected size of the mutant protein. The C-terminal SPTAN1 antibody is downstream of the out-of-frame indel mutation and is not expected to identify the mutant allele. **E)** Fold change in the number of mutant RNA transcripts from deep-targeted RNA sequencing of heterozygous mutated genes in NCI-H358 and LS180 xenografts treated by oral gavage with 60mg/kg LY3023414 assayed 16 hours post-treatment. Student’s t-test for target genes are all p < 0.05, while the null hypothesis holds for *RNF43* (common SNP).

The whole transcriptome sequencing data was confirmed by targeted deep RNA sequencing of two naturally occurring heterozygous truncating mutations from each cell line. Increased expression of the truncated mutant alleles relative to the wild-type alleles of the *EXOC1* and *SPTAN1* genes was observed in NCI-H358 cells after treatment with LY3023414 (Figure 2B left). Likewise, increased expression of *HNRNPA3* and *LRPPRC* mutant transcripts was observed in LS180 after LY3023414 treatment (Figure 2B right). We noted no significant effects on two coding region heterozygous single nucleotide polymorphisms (SNP’s) after LY3023414 treatment in the genes *RNF43* or *DROSHA* in either cell line (Figure 2B). To substantiate these transcriptomic effects at the protein level, we treated NCI-H358 cells for 24 hours with LY3023414. Western blotting revealed truncated mutant protein for both EXOC1 and SPTAN1 after treatment with LY3023414, whereas the full-length protein levels remained present and unchanged in both the treated and control (DMSO) samples (Figure 2C and 2D left)[41]. An antibody that recognizes the C-terminus of SPTAN1, encoded downstream of the frameshift mutation, did not detect mutant protein, as expected (Figure 2D right).

LY3023414 was originally developed as a phosphatidylinositol 3-kinase (*PI3K*) inhibitor with activity against AKT Serine/Threonine Kinase 1 (*AKT1*) and Mammalian Target of Rapamycin (*mTOR*)[42]. It was tested in a number of clinical trials and has a well-known pharmacokinetic profile *in vivo*[43-47]. To determine whether LY3023414 affects NMD *in vivo*, we established xenograft tumors of both NCI-H358 and LS180 in nude mice. Treatment of these mice with a single oral dose of 60 mg/kg LY3023414 led to a significant increase in the expression of mutant RNA transcripts relative to wild-type transcripts 16 hours later (Figure 2E). The *RNF43* gene, which harbors a coding region heterozygous SNP, served as a control (Figure 2E). Severe drug-associated toxicity, including weight loss bordering on cachexia and near total inactivity, precluded the longer-term dosing required for anti-tumorigenic effects of LY3023414 in both BALB/c and C57BL/6N immunocompetent mouse strains.

### The kinase SMG1 is the target for NMD inhibition by LY3023414

To investigate the mechanism of NMD inhibition by LY3023414, we evaluated the six kinases with the highest reported inhibition by LY3023414[42]. siRNA-mediated knockdown of each of these kinases in NCI-H358 and LS180 cancer cell lines was performed for this purpose (Figure S9). Only knockdown of *SMG1* resulted in significant changes in the amount of truncating mutant transcript relative to wild-type transcript in all four genes evaluated (Figure 3A top). *RNF43* and *DROSHA* contain heterozygous coding region SNP’s and served as controls in these experiments and as expected showed no changes despite siRNA treatment (Figure 3A bottom).

**Figure 3:**
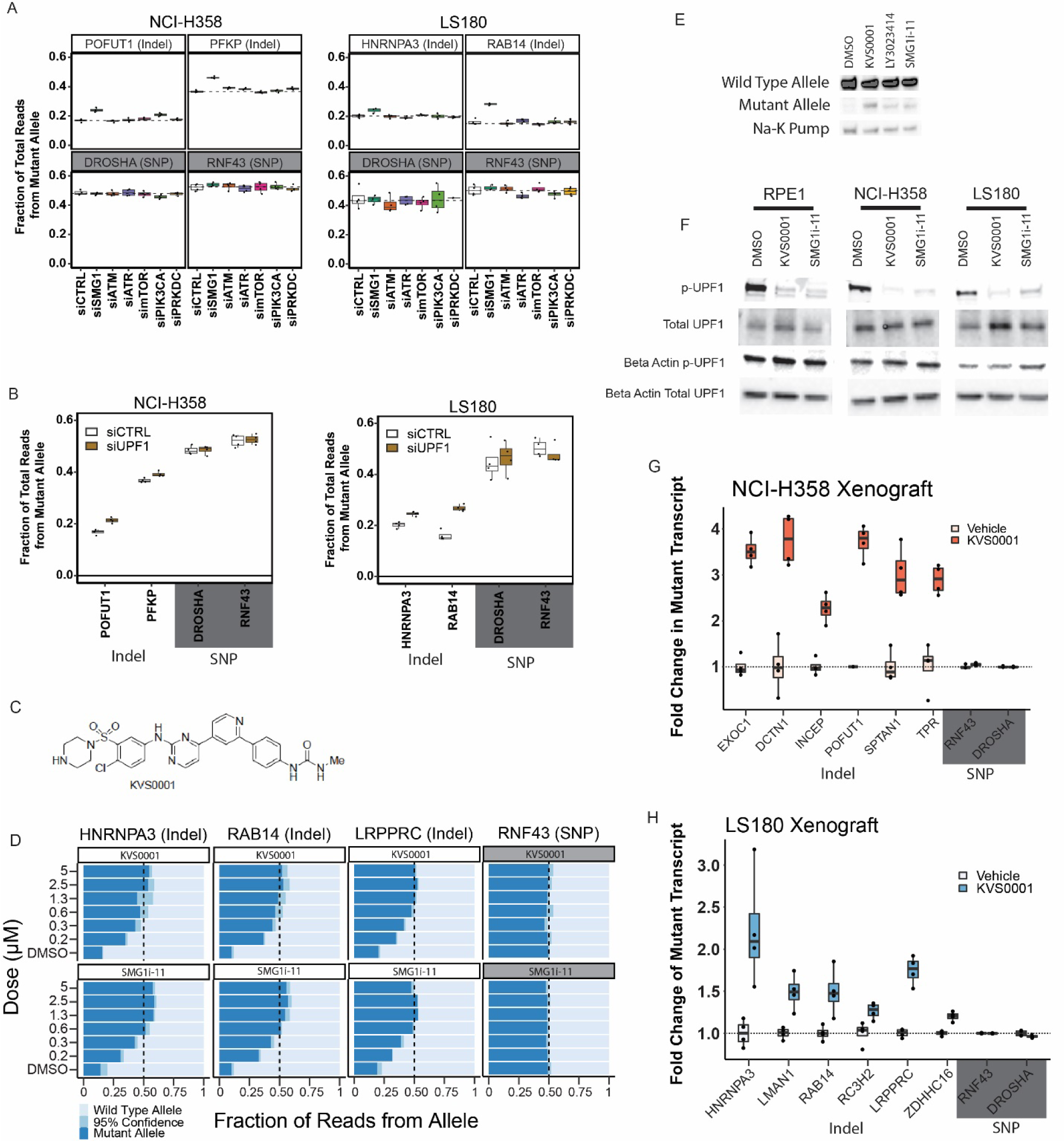
Novel NMD inhibitor KVS0001 is SMG1 specific and induces expression of NMD targeted genes *in vitro* and *in vivo*. **A)** Fraction of mutant allele transcripts in genes with heterozygous indels previously established in this study as sensitive to NMD inhibition. Results show mutant levels after siRNA treatment targeting kinases inhibited by LY3023414. *RNF43* and *DROSHA* are common heterozygous SNPs (shaded gray) and serve as negative controls. **B)** Fraction of mutant allele transcripts in genes with truncating mutations known to be sensitive to NMD inhibition after siRNA treatment with siUPF1 or non-targeting siRNA. Data from deep-targeted RNA-sequencing. **C)** Structure of novel NMD inhibitor KVS0001. **D)** Targeted RNA-sequencing on three genes with heterozygous, out-of-frame, indel mutations in LS180 cancer cells treated in a dose-response with KVS0001 or SMG1i-11. *RNF43* serves as a control (common heterozygous SNP) and the mutant allele refers to the non-reference genome allele. **E)** Western blot of EXOC1 protein in NCI-H358 cells treated with 5µM novel inhibitor KVS0001, LY3023414 or SMG1i-11 for 24 hours. **F)** Western blot of phosphorylated UPF1 on three cell lines treated with 5µM KVS0001, SMG1i-11, or DMSO. Note that total UPF1 and p-UPF1 were run on different gels, loading controls correspond to indicated gel. **G)** Fold change in the number of mutant allele transcripts measured by targeted RNA-Seq in genes containing heterozygous out-of-frame indel mutations in NCI-H358 or **H)** LS180 subcutaneous xenografts in bilateral flanks of nude mice. Mice were treated once with IP injection of vehicle or 30mg/kg KVS0001 and tumors harvested 16hrs post IP treatment. All genes shown contain heterozygous out-of-frame truncating mutations except *RNF43* and *DROSHA* which serve as controls (contain heterozygous SNP’s).

*SMG1* is known to regulate the NMD pathway by activating *UPF1*, an enzyme with RNA helicase and ATPase activity[48, 49]. We therefore knocked down *UPF1* with siRNA and found that it restored expression of the NMD-downregulated transcripts, at levels similar to those observed after the knock-down of *SMG1* (Figure 3B). Additionally, treatment of NCI-H358 and LS180 cancer cell lines with a previously reported *SMG1* specific small molecule inhibitor, SMG1i-11 resulted in specific increases in the transcripts and proteins from genes with truncating mutations, just as did LY3023414 (Figures S10 and S11)[32]. Although SMG1i-11 displayed considerable *SMG1* specificity and marked inhibition of NMD, it was highly insoluble. While we had no difficulty getting it into solution in DMSO for *in vitro* work, we were unable to find a vehicle to administer it *in vivo*, despite numerous attempts at various formulations and administration routes. This may explain why SMG1i-11 has been demonstrated to be an effective *SMG1* inhibitor *in vitro*, but no peer-reviewed reports of its *in vivo* activity have been reported to date[32, 50, 51].

### Development of an improved NMD inhibitor targeting SMG1

Although we were unable to secure a viable lead compound, the HTS did identify *SMG1* as an ideal target to disrupt the nonsense-mediated decay pathway. We sought to develop a new SMG1 inhibitor based on the cryo-electron microscopy structure of the binding pocket of SMG1[52, 53]. We attempted the synthesis of eleven compounds (KVS0001 to KVS0011) and tested for bioavailability and preservation of target specificity. Among these, KVS0001 stood out due to its solubility while preserving SMG1 inhibitory activity (Table S2, Figures 3C and S12). Mass spectrometry-based assays showed that KVS0001 inhibits SMG1 protein more than any of the other 246 protein or lipid kinases tested at concentrations from 10nM to 1µM (Table S4 and Figure S13)[54]. Noteworthy off-target kinase inhibition was not observed until doses of 1µM and above. Experiments using NCI-H358 and LS180 cells showed that KVS0001 is bioactive in the nanomolar range and subverts the NMD-mediated down-regulation of truncating mutant transcripts and proteins (Figures 3D, 3E, S14, S15). Inhibition at concentrations as low as 600nM led to equal expression of wild-type and mutant transcripts, suggesting near total blockade of the NMD pathway (Figures 3D, S14). Western blotting showed that KVS0001 substantially decreases the amount of phosphorylated UPF1, the downstream target of SMG1 kinase activity, in three different cell lines (Figure 3F). Finally, KVS0001 treatment of NCI-H358 and LS180-derived xenograft tumors in nude mice resulted in significant increases in transcript levels in each of six tested endogenous genes with truncating mutations, while having had no measurable effects on genes containing heterozygous coding region SNP’s (Figures 3G and 3H).

### NMD inhibition with KVS0001 causes MHC class I display of hidden neoantigens

Cancer cells may evade immune surveillance by downregulating genes with truncating mutations as a result of NMD[19, 55]. Indeed, previous studies with non-specific or toxic NMD inhibitors have shown an increase in selected antigens from tumor-specific mutations when NMD is inhibited[56]. We therefore investigated whether KVS0001 could similarly alter the cell surface presentation of proteins from genes harboring truncating mutations in NCI-H358 and LS180 cells. Using quantitative HPLC-mass spectrometry (MS) we observed a striking (45 to 90 fold) increase in the *EXOC1* and *RAB14* derived neoantigens, and a significant (2 fold) increase in the *ZDHHC16* derived neoantigen (Figures 4A, 4B, S16, S17)[57]. This is consistent with the re-expression of mutant transcript and protein shown previously in this study (Figures 3D and 3E). These three peptides were chosen based on an *in-silico* review of potentially presented peptides and a preliminary experiment that looked at qualitative (present or not present) presentation of the predicted peptides[58, 59].

**Figure 4:**
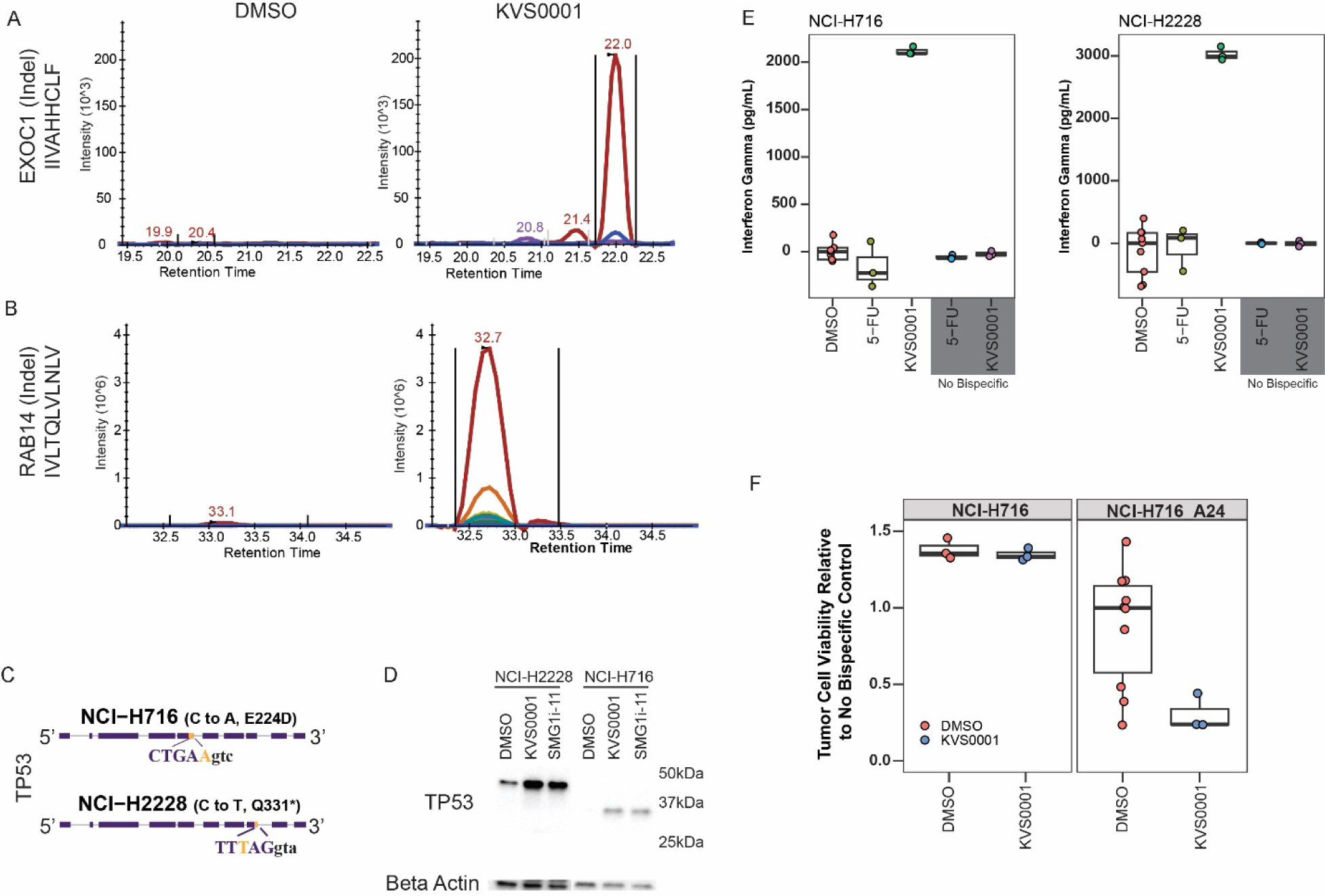
KVS0001 treatment induces targetable cell surface presentation of peptides known to be downregulated by NMD. **A)** MHC class I HLA presentation of mutant specific peptide sequences from NCI-H358 and **B)** LS180 cells by quantitative HPLC-Mass Spectrometry treated with DMSO or 5µM KVS0001. The gene name, type of mutation (in parenthesis) and presented peptide are shown on the y-axis for each gene. Colors indicate different ions. **C)** *TP53* gene structure and mutant DNA sequence for NCI-H716 and NCI-H2228 cancer cell lines, both contain a homozygous splice site mutation in *TP53*. Capital letters represent exonic sequence; lowercase letters represent intronic sequence. DNA mutation reflected by gold bases. **D)** Western blot against TP53 in the presence or absence of 5µM NMD inhibitor in NCI-H716_A24 and NCI-H2228 cell lines. NCI-H2228 has an expected size of 46.6 kDa and NCI-H716 of 34.7 kDa. **E)** IFN-γ levels over baseline based on ELISA in a co-culture assay with NCI-H716_A24 and NCI-H2228 cells, 1.25µM NMD inhibitor, human CD3+ T-cells, and bispecific antibody for TP53 and CD3. Chemotherapy (5-Fluorouracil) is shown as a control. **F)** Cell killing based on luciferase levels in a co-culture assay in NCI-H716 cells with and without A24 expression, treated with TP53-CD3 bispecific antibody, 1.25µM NMD inhibitor and human CD3+ T-cells.

### Targetable peptide presentation occurs with NMD inhibition by KVS0001

To test whether cancer cell neoantigens presented as a result of NMD inhibition could be targeted, we evaluated two cancer cell lines, NCI-H716 and NCI-H2228 (Table S1). Both lines contain homozygous truncating mutations in *TP53* which produce transcripts down-regulated by NMD, with each exhibiting RNA levels less than 20% of the median level of expression among 675 cancer cell lines (Figures 4C and S18)[60, 61]. Treatment with KVS0001 increased the expression of TP53 in both lines, while the commonly used therapeutic agents 5-fluorouracil and etoposide did not affect TP53 protein abundance (Figures 4D and S19).

To determine whether this disruption of NMD repression is targetable by T-cells, we developed a bispecific antibody (KVS-BI043) that recognizes a peptide-HLA complex on one end and CD3 on the other end. CD3 is expressed only on T-cells, and this bispecific antibody functions as a T-cell engager, linking target cells to cytotoxic T-cells, which then kill the targets[62, 63]. KVS-BI043 recognizes a ten amino acid peptide (residues 125 to 134 of TP53) bound to HLA-A24. NCI-H2228 naturally express A24 whereas NCI-H716 cells were engineered to express A24 using a retrovirus (NCI-H716_A24). Treatment of NCI-H716_A24 or NCI-H2228 cells with KVS0001 and the KVS-BI043 bispecific antibody in the presence of normal T-cells caused a significant increase in interferon (IFN-γ) release (Figure 4E). We observed no changes in IFN-γ levels in the absence of KVS-BI043 or normal T-cells (Figure 4E grey boxed lanes). Most importantly, treatment of NCI-H716_A24 with KVS0001 in the presence of KVS-BI043 and T-cells led to significant killing of the target cancer cells, which was also not observed in the absence of T-cells or the absence of HLA-24 in the target cells (Figure 4F). NCI-H2228 was assessed for killing but expressed too much TP53 at baseline (Figure 4D left most lane) and thus displayed substantial killing even in the DMSO controls.

### Tumor growth is slowed in mice treated with KVS0001

Finally, we investigated whether KVS0001 could impact tumor growth in syngeneic models in which the native immune system might play a role (Table S1). For this purpose, we first used murine RENCA (renal cancer) and LLC (lung cancer) as they are known to have a relatively large number of out-of-frame indel mutations (Table S5). Although murine and human SMG1 are highly related (98% at the amino acid level), it was important to show that KVS0001 could actually inhibit NMD in murine cells.

For this purpose, we tested eight genes in LLC, and four in RENCA, which contained out-of-frame indel mutations potentially targeted by NMD. We also assessed the expression of three genes without any mutations known to have their normal expression controlled by NMD[64]. We found that six of the twelve truncating mutation-containing genes and five of the six expression controlled by NMD genes had significantly increased RNA following *in vitro* treatment with KVS0001 (Figure 5A and 5B).

**Figure 5:**
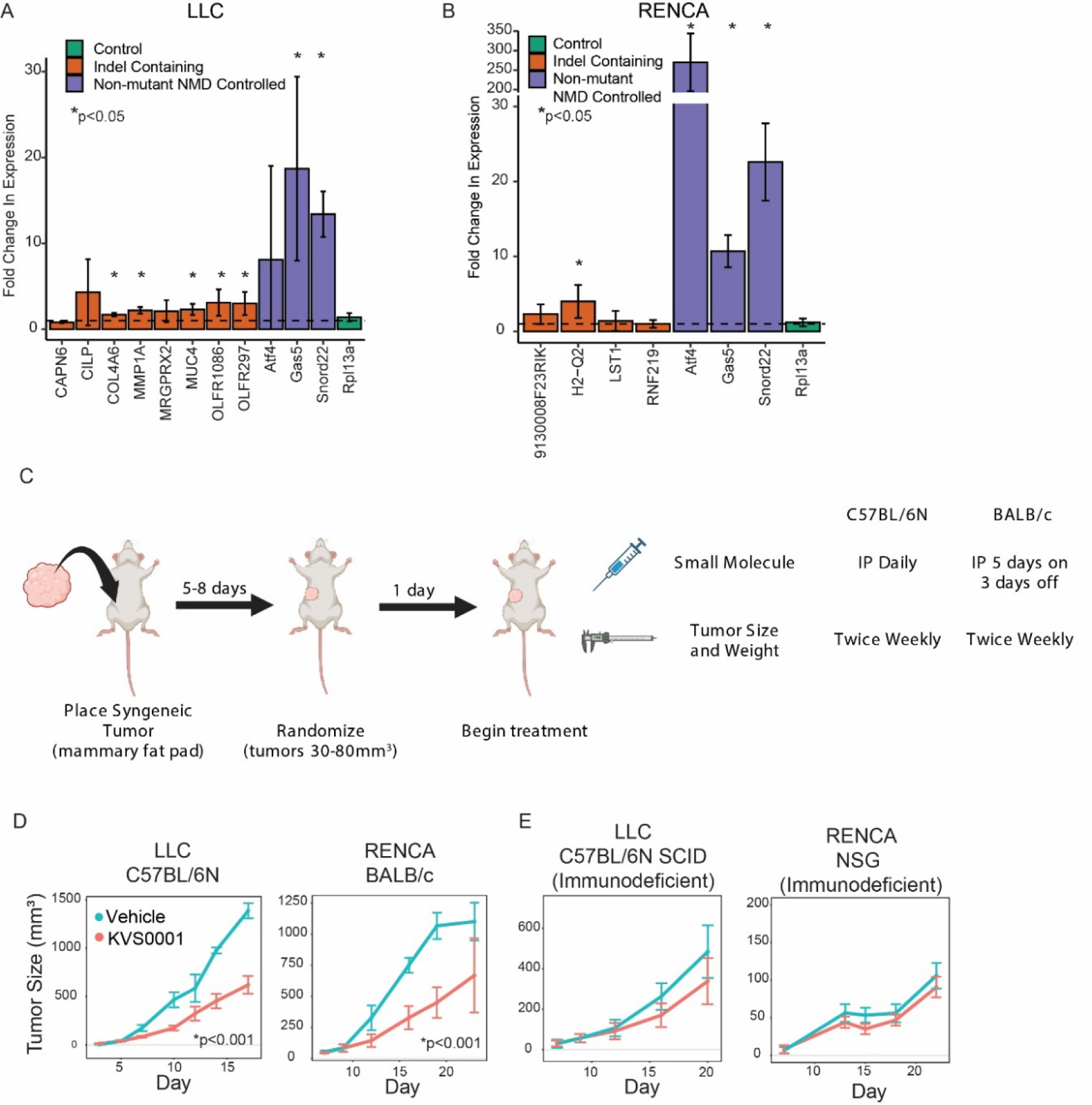
*In vivo* treatment of murine tumors with KVS0001 yield differential tumor growth compared with vehicle treatment. **A)** Fold change in RNA transcript levels in LLC or **B)** RENCA cells treated in-vitro with 5µM of NMD inhibitor KVS0001 or DMSO. Orange bars indicate genes with homozygous indel mutations potentially targeted by NMD. Purple bars show genes with no mutations but that are known to have their normal transcription levels controlled by NMD. Green bar is a control gene that should not change with treatment. The dotted line shows relative expression of DMSO treatment (equal to 1). **C)** Treatment schedule for syngeneic tumor mouse experiments. **D)** Average tumor size of LLC (left) and RENCA (right) syngeneic tumors in immune-competent mice (n=8) treated with 30 mg/kg KVS0001 or vehicle control. Difference is statistically significant after day 10 based on one-way ANOVA with Dunnett’s test p<0.001 (p<0.05 for day 23 RENCA data point) for both tumors tested. **E)** Average tumor size of LLC (left) and RENCA (right) in immunodeficient mice (n=8) treated with 30 mg/kg KVS0001 or vehicle control. Error bars show 95% confidence intervals in all plots.

We then implanted LLC and RENCA cancer cells in the mammary fat pad of C57BL/6N and BALB/c mice respectively and treated with KVS0001 or vehicle control (Figure 5C). The dose of KVS0001 was based on experiments showing that the maximum solubility limit was reached around 2 to 3mg/mL, leading to a maximum single dose of 30mg/kg per mouse per treatment. At this dose, the only toxicity noted was transient weight loss (Figure S20), and no other pathology was observed. Both tumor types experienced significant slowing of tumor growth (Figure 5D). However, when the same tumors were implanted in immunocompromised mice, there was no statistically significant difference in tumor growth between mice treated with KVS0001 or vehicle control (Figure 5E).

We evaluated a total of seven syngeneic mouse cancer cell lines. Four contained a relatively high number of truncating indel mutations, while the remaining three contained a moderate or low number. None of the three models with moderate or low numbers of truncating mutations responded to KVS0001 (EMT6, B16-F10, M3 Melanoma) (Figure S21). Of the four models with a higher number of truncating mutations, two (LLC and RENCA) responded in a statistically significant fashion while the other two failed to reach significance (CT26, MC38) (Figures 5D and S21).

## Discussion

We report the design and execution of a high throughput screen to identify small molecule inhibitors of the nonsense-mediated decay pathway. The method employs a ratiometric output, allowing quantitative and controlled determination of RNA expression changes associated with NMD. Unlike previous NMD screens, this assay is directed at NMD as a process rather than as a way to identify compounds for treating specific mutations related to a disease state, such as cystic fibrosis or β-thalassemia[28, 30, 65]. The genetically modified isogenic cell lines employed here are uniquely suited for this approach, producing a high signal-to-noise ratio. Follow-up experiments confirmed a low false positive rate in the screen: four of the eight hits re-expressed transcripts from genes with truncating mutations in a dose-responsive manner when re-tested. Note that our assay allows for the detection of NMD inhibitors regardless of effects on protein translation by virtue of the comparison of wild-type to mutant transcripts for each queried gene. This is an important feature, as many previously reported inhibitors of NMD, such as emetine and anisomycin, rely on inhibition of protein translation and thus are unlikely to be optimal for specific restoration of NMD-targeted RNA and protein expression[35, 38, 39, 66]. During the review process it was also noted that different NMD inhibitors caused re-expression of different knockout clones to different extents (see Figure 1B and S8 as an example). It is not clear to us why this occurred, but we speculate that it may be related to differences in the mechanism of action (i.e., protein synthesis inhibition based NMD effects versus targeted NMD pathway inhibition as one example).

The optimal inhibitor of NMD found in our screen was LY3023414, and we suspected that this compound may inhibit the kinase *SMG1* based on previously published kinase activity data with this small molecule[42]. *SMG1* acts as the gatekeeper of *UPF1* activity through its phosphorylation at multiple sites[48]. Unfortunately, issues likely related to off-target (pan-kinase) toxicity at the doses required for NMD inhibition prevented LY3023414 from being utilized in further *in vivo* work. While there are previously described SMG1 inhibitors, all have undesirable off-target or biophysical characteristics complicating their use as therapeutic agents[28-31, 39, 56, 67]. For example, SMG1i-11 was previously identified as a potent and specific SMG1 inhibitor[32]. However, in our hands, it was not soluble at concentrations required for in vivo work in a suitable vehicle for mouse administration.

Another unrelated compound has shown successful *in vivo* re-expression of mutant RNA transcripts, but the effect is reported as less than a two-fold change, compared to the three to four-fold demonstrated here by KVS0001, and little work has been done to develop the compound further in subsequent studies[68].

Despite these chemistry related challenges, our screen offered an unbiased assessment of the best targets for NMD pathway disruption, and thus we concluded that SMG1 was an ideal protein to inhibit for this purpose. In light of this, we attempted to improve the biophysical properties of SMG1i-11 through the development of KVS0001. As reported in this study, KVS0001 was a specific inhibitor of SMG1 that was soluble and could easily be administered to mice. Treatment of KVS0001 not only increased levels of transcripts and proteins from genes with truncating mutations, but also increased the levels of corresponding peptide bound-HLA complexes on the cell surfaces. We developed a new bispecific antibody, KVS-B043, which could recognize these pHLA complexes when the truncating mutations were in the *TP53* gene and destroy cancer cells harboring such mutations. In syngeneic mouse models, KVS0001 slowed the growth of some tumors containing multiple truncating mutations, but only when the mice were immunocompetent. This dependence on an intact immune system supports the idea that the slowed growth of the treated tumors was due to recognition of neoantigens.

Despite these encouraging data, the potential utility of KVS0001 for therapeutic purposes remains speculative, for several reasons. First, we do not understand why KVS0001 (and other NMD inhibitors) increase the expression of many, but not all, genes with truncating mutations. We also do not systematically address the changes to splicing that are undoubtedly occurring when NMD is disrupted in either normal or tumor tissue. More research on the biochemistry of NMD will undoubtedly be required for such understanding, and perhaps the tools described here may facilitate that effort. Second, treatment with KVS0001 reduced the growth of some but not all syngeneic tumor types tested in mice. There appeared to be a correlation between the ability of KVS0001 to inhibit growth and the number of truncating mutations in the tumor. Likewise, recent evidence has been presented showing that cells deficient in known splicing factors are sensitive to subsequent NMD inhibition[69]. This situation is reminiscent of that encountered with immune checkpoint inhibitors, in which there is a positive but imperfect correlation with loss of microsatellite instability (MSI) related proteins and tumor mutation burden[8, 12]. Undoubtedly, the ability of the immune system to react to neoantigens is related to their quality as well as their quantity, with quality defined as the ability of the neoantigen to be bound to the host’s particular MHC constitution and initiate an immune response[70, 71]. Third, treatment with KVS0001 led to decreased tumor growth in immunocompetent mice, but did not lead to tumor regressions of the type mandated by RECIST criteria in human clinical trials[72, 73]. Whether this failure to cure tumors in mice results from the contrived nature of the models used – injection of large numbers of rapidly growing cancer cells into animals with relatively little time to react prior to their demise – complicates the interpretation of many pre-clinical immunotherapeutic approaches.

Finally, KVS0001, though it was soluble and could be administered to animals for at least a month, was not entirely non-toxic, causing transient weight loss in some of the mice. Previous work has raised concerns that the importance of NMD in development and normal gene expression make it a difficult pathway to safely disrupt[21, 74-77]. The limited impact on body weight or observed physical activity of mice undergoing KVS0001 therapy at therapeutically relevant doses suggests that NMD inhibition may have acceptable toxicity and be tolerated in developed animals. Another example of this lies in the natural genetic variants of NMD components, which convey variability in the efficiency of NMD between humans, and lends itself to supporting that knockdown of this pathway may be tolerable[78-80]. Studies in mice also support this observation, as post-development knockdown of UPF1 displayed minimal phenotype change[64]. The broad use of NMD in normal development and growth, coupled with the observations here, suggests future work with inhibitors of this pathway should continue with a close eye for on-target off-tissue toxicity.

Though there is much work to be done, we hope that the tools, approaches, and compounds described here will facilitate that work. Should the administration of KVS0001 or a related compound prove non-toxic and well-tolerated in humans, we end with a speculation about the possibility of KVS0001 being used as a preventative rather than as a therapeutic agent. Its potential use to prevent the onset of symptoms in pediatric syndromes caused by germline truncating mutations is obvious[17, 81]. Less obvious is the potential for it to prophylactically reduce cancer incidence in patients with hereditary non-polyposis colorectal cancer. These patients inherit heterozygous mutations of a mismatch repair gene, and they do not develop tumors until biallelic mutations of that mismatch repair gene are acquired in a rare stem cell during the second or third decade of life. Similar speculations can be made about the potential of KVS0001 to be used to prevent cancer initiation or progression in patients with other inherited mutations in repair genes, or in individuals exposed to high levels of exogenous mutagens.

## Acknowledgements

The authors thank Dr. Qing Wang from Complete Omics Inc. for his help and advice with the neoantigen presentation by mass spectrometry studies. We would also like to thank Dr. Xiao Chen from Ascendex LLC. for helpful suggestions during medicinal chemistry experiments. Biophysical predictions for small molecules was done using Molinspiration Cheminformatics free web services, https://www.molinspiration.com, Slovensky Grob, Slovakia, accessed 9/2023. This work was supported by Oncology Core CA 06973 (B.V., K.W.K., N.P.); The Virginia and D.K. Ludwig Fund for Cancer Research (B.V., K.W.K., N.P., C. B.); The Sol Goldman Sequencing Facility at Johns Hopkins (B.V.); NIH Cancer Center Support Grant P30 CA006973. SP was supported by NCI Grant K08CA270403, the Leukemia Lymphoma Society Translation Research Program award, the American Society of Hematology Scholar award, and the Swim Across America Translational Cancer Research Award.

## Author contributions

A.L.C, K.W.K., and N.W. conceptualized the study. A.L.C, S.S., B.V., N.P., C.B., K.G., S.Z., K.W.K., and N.W. developed the methodology. A.L.C., S.S., L.D., E.W., B.P., B.S.L., S.P., E.H., M.P., and N.W. conducted the experiments. A.L.C., S.S., J.D.C., B.S.L., and N.W. performed the formal data analysis. A.L.C, N.W., and K.W.K. wrote the paper.

## Supplemental Materials

### STAR methods

#### KEY RESOURCE TABLE

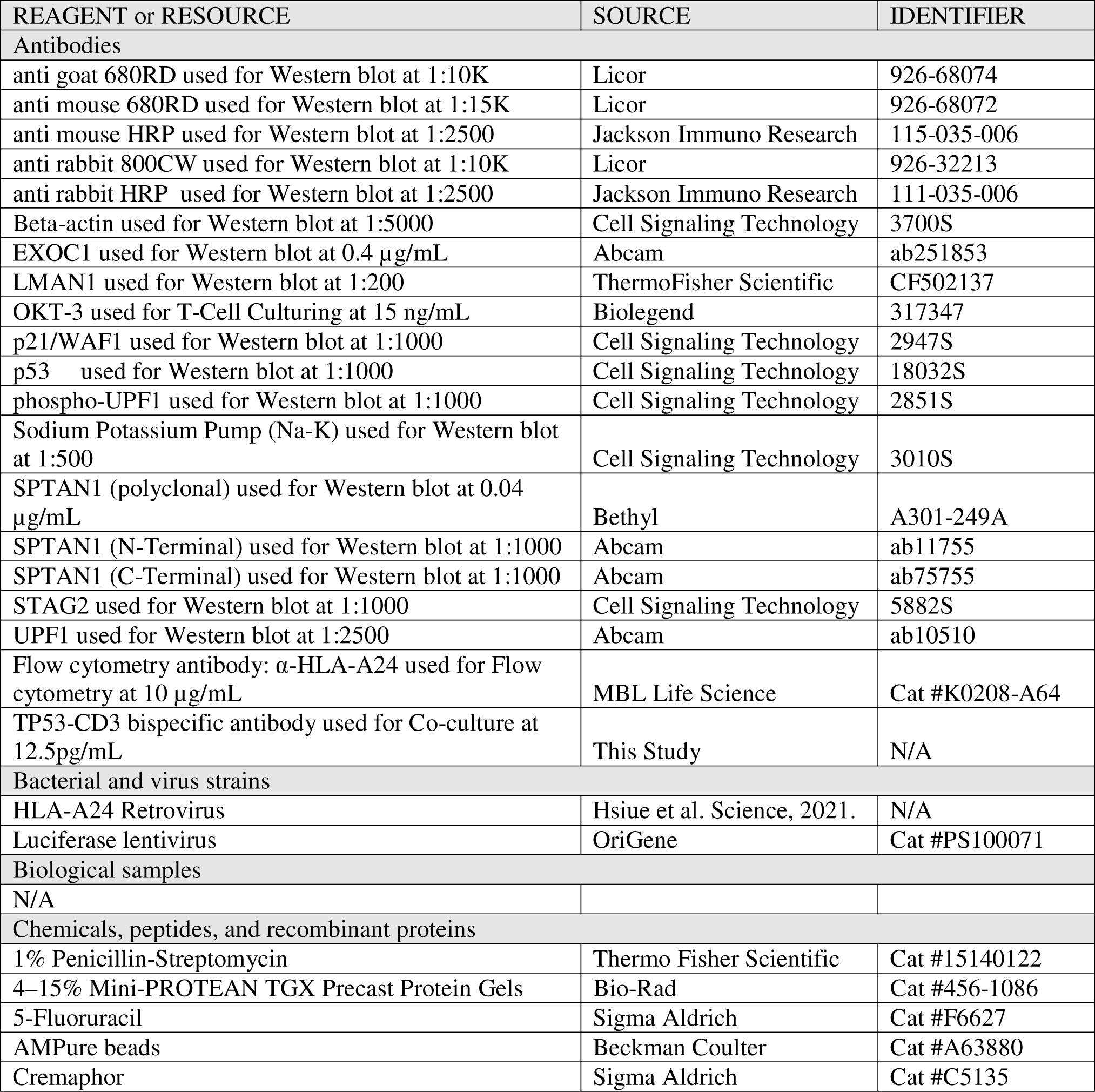
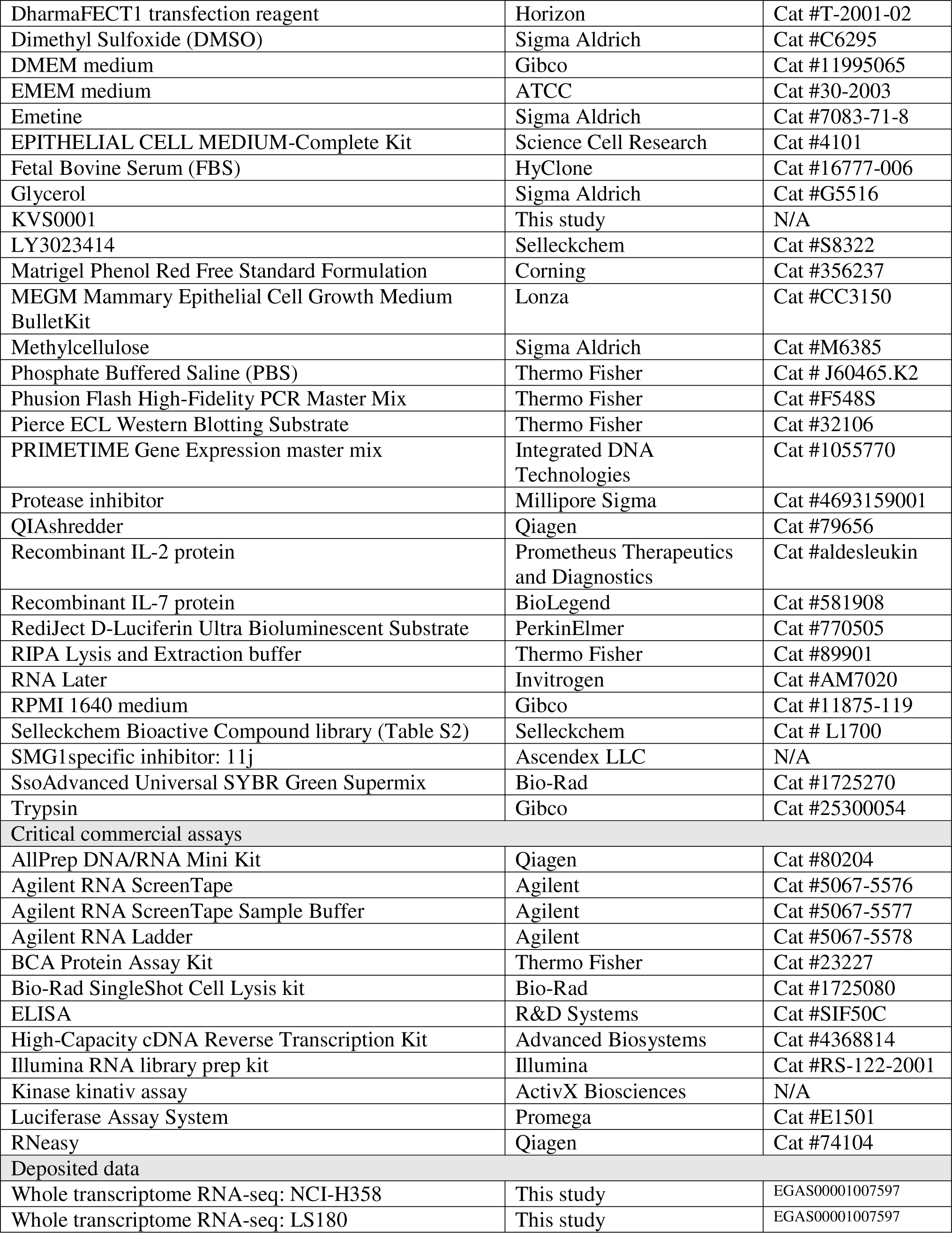
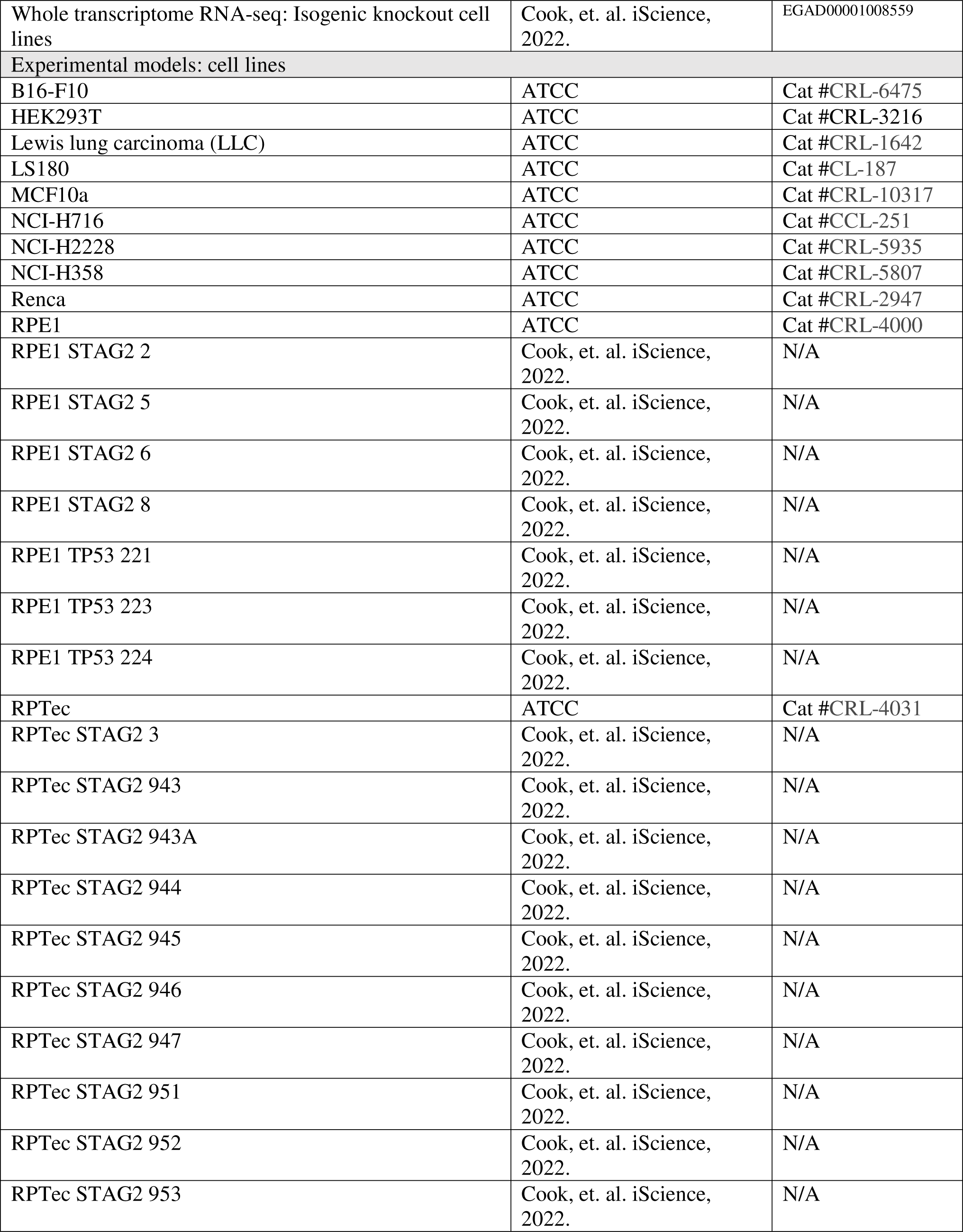
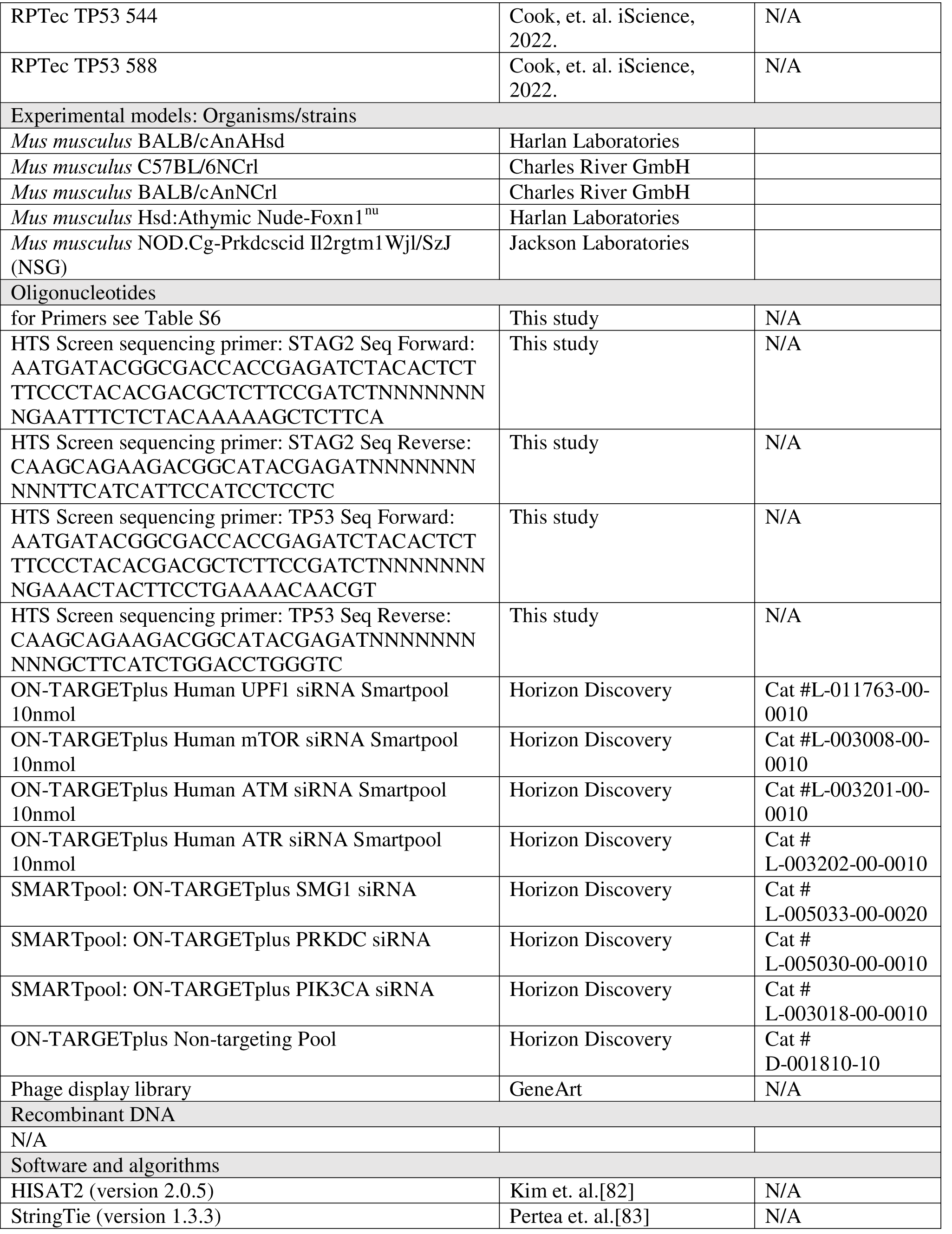
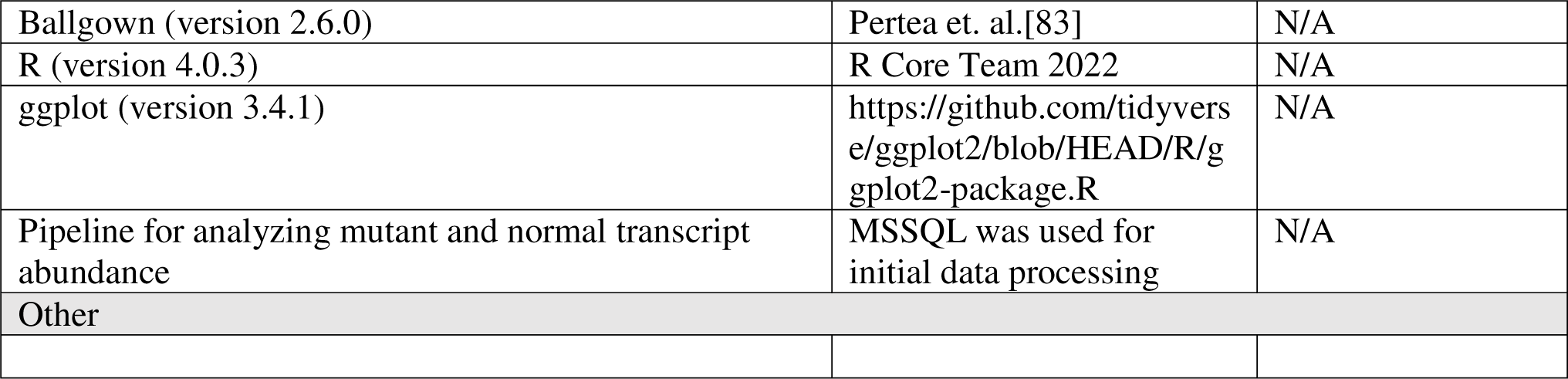

#### RESOURCE AVAILABILITY

##### Contact for Reagent and Resource Sharing

Further information and requests for resources and reagents should be directed to and will be fulfilled by the lead contact, Nicolas Wyhs. Isogenic knockout cell lines are available through The Genetic Resources Core Facility at Johns Hopkins School of Medicine (, Maryland, USA). KVS0001 generated in this study will be made available on request if available, but availability may be limited and we may require a payment and/or a completed materials transfer agreement if there is potential for commercial application.

##### Data availability

- Whole transcriptome RNA-seq raw and processed data have been deposited at GEO and are publicly available as of the date of publication. Accession numbers are listed in the Key Resource Table.
- Full western blot images have been deposited at Mendeley (Mendeley Data, V1, doi: 10.17632/n5s2z393z6.1) and are publicly available as of the date of publication.
- Any additional information required to reanalyze the data reported in this paper is available from the lead contact upon request.

### EXPERIMENTAL MODELS AND SUBJECT DETAILS

#### Cell lines

NCI-H716, NCI-H2228, HEK293, NCI-H358, RPE1, MCF10a, RPTec, LLC, RENCA, B16-F10 and LS180 cells were purchased from The American Type Culture Collection (Virginia, USA). RPE1 STAG2 2, RPE1 STAG2 5, RPE1 STAG2 6, RPE1 STAG2 8, RPTec STAG2 3, RPTec STAG2 943, RPTec STAG2 943A, RPTec STAG2 944, RPTec STAG2 945, RPTec STAG2 946, RPTec STAG2 947, RPTec STAG2 951, RPTec STAG2 952, RPTec STAG2 953, RPE1 TP53 221, RPE1 TP53 223, RPE1 TP53 224, RPTec TP53 54, RPTec TP53 588, RPTec TP53 544 isogenic knockout cell lines were generated and grown as previously described (Cook et al, iScience 2022)[33]. NCI-H358, NCI-H2228, NCI-H716, RENCA and RPE1 cells were grown in RPMI 1640 Medium (Gibco, California, USA, Cat #11875-119) supplemented with 10% Fetal Bovine Serum (FBS) (HyClone, Utah, USA, Cat #16777-006). LS180 cells were grown in EMEM (ATCC, Virginia, USA, Cat #30-2003) supplemented with 10% FBS. LLC, B16-F10 and HEK293 was grown in DMEM (Gibco, USA, Cat #11995065) supplemented with 10% FBS. RPTec cells were grown in EPITHELIAL CELL MEDIUM-Complete Kit (Science Cell Research, California, USA, Cat #4101). MCF10a cells were grown in MEGM Mammary Epithelial Cell Growth Medium BulletKit (Lonza, USA, Cat #CC3150). *In vitro*, all cells were grown at 37°C with 5% CO_2_.Mycoplasma testing was performed by The Genetic Resources Core Facility at Johns Hopkins School of Medicine (Maryland, USA).

#### Quantitative real-time PCR (qPCR)

RNA was obtained from cells using a Qiagen RNeasy Kit (Qiagen, Maryland, USA, Cat #74104) per the manufacturer’s instruction. Reverse transcription of RNA to cDNA was performed using a High-Capacity cDNA Reverse Transcription Kit (Applied Biosystems, USA, Cat #4368814) per manufacturer’s instructions. Unless otherwise specified, qPCR reactions were set up using SsoAdvanced Universal SYBR Green Supermix (Bio-Rad, USA, Cat #1725270) following manufacturer’s instructions and performed on a QuantStudio 3 (Applied Biosystems, USA) with manufacturer’s recommended plates and plate covers. PCR thermocycling conditions were as follows: 2 min at 50°C, 2 min at 95°C, 35 cycles of 10 sec at 95°C, 10 sec at 60°C, 30 sec at 72°C. Finally a melt curve was performed starting at 50°C and ending at 95°C with 5-second incubations for imaging. For TP53β analysis, qPCR reactions were set up using PRIMETIME™ Gene Expression master mix (Integrated DNA Technologies, USA, Cat #1055770) and a probe with FAM reporter and TAMRA quencher (Integrated DNA Technologies, USA) following the manufacturer’s instructions and concentrations. PCR thermocycling conditions were as follows: 3 min at 50°C, 10min at 95°C, 40 cycles of 10 sec at 95°C, 30 sec at 60°C. Primers used for all qPCR can be found in Table S6.

#### Small molecule compounds

SMG1 specific inhibitor SMG1i11 (11j)[32] and KVS0001 were synthesized by Ascendex LLC (Pennsylvania, USA). The synthesis scheme for KVS0001 is located below. Emetine was obtained from Sigma Aldrich (Cat #7083-71-8). All other small molecule hits from the screen were purchased from Selleckchem (Texas, USA).

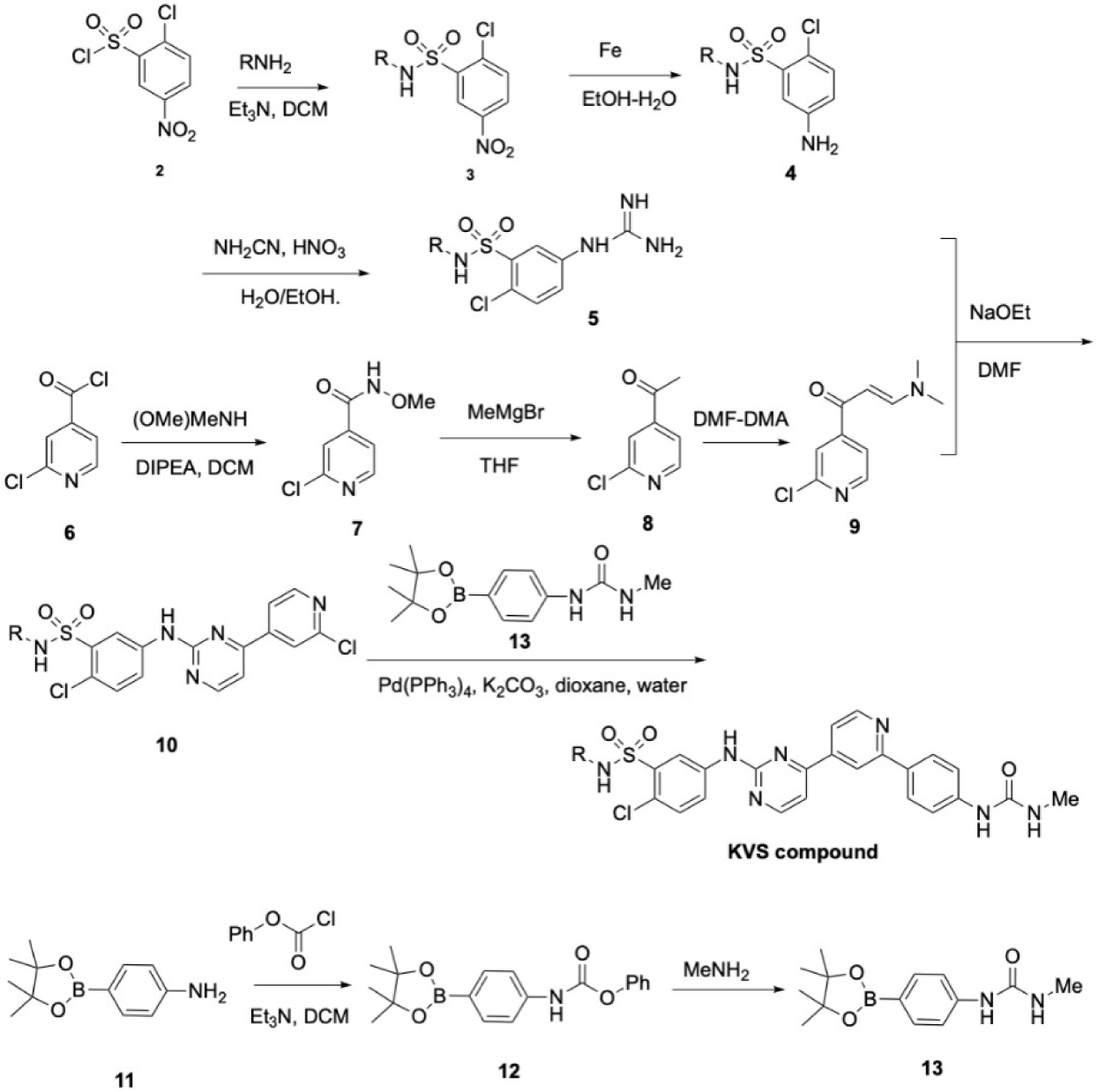

#### NMD screen

We screened the Selleckchem Bioactive Compound library, a 2,658 compound library (Selleckchem, Texas, USA, Cat #L1700). The library was distributed over 33 x 96-well tissue culture plates with each plate containing library compounds, 2 NMD positive controls (emetine, 12 μg/mL) and 6 NMD negative controls (DMSO). We mixed the three cell lines, RPE1 STAG2 2, RPE1 STAG2 8, and RPE1 TP53 221, in equal ratio and dosed with the compound library at 10 μM for 16 hours. Cell pools were harvested by washing plates 2x with PBS and frozen. RNA extraction and reverse transcriptase cDNA conversion were performed as described in *Targeted RNA-seq.* Quality control was performed by qPCR of one NMD positive and one NMD negative control well from each plate to confirm that increased expression of the positive control (emetine) wells was observed. All samples were prepared for sequencing by amplifying cDNA using primers containing both a plate (forward primer) and well (reverse primer) barcode which amplifies the knockout cell line specific mutation for STAG2 and TP53 (See Table S6). Primers were obtained from Integrated DNA Technologies (Iowa, USA). PCR was performed using Phusion Flash High-Fidelity PCR Master Mix (Thermo Fisher, USA, Cat #F548S) for 1 min at 98 °C, 29 cycles of 10 sec at 98 °C, 15 sec at 64 °C, 15 sec at 72 °C, and 5 min at 72 °C. Samples were then well barcoded using a similar PCR setup for 2-4 cycles. Samples were then pooled, cleaned up with AMPure beads (Beckman Coulter, California, USA, Cat #A63880), and sequenced on an Illumina HiSeq2500 using manufacturer’s instructions (150 cycle single read) for an average of 56,753 reads per well. The screen was scored by calculating the number of sequencing reads matching each mutant and reference transcript and calculating a Mutant Allele Fraction (MAF) correcting for the number of cell lines and heterozygous mutations.

NMD positive and negative control data were pooled, respectively, and averaged across all plates to determine the hit threshold of MAF > 0.46. This value, 5 standard deviations above the mean for the DMSO controls, was chosen to ensure no false positives in the DMSO controls were observed. A compound was considered a hit if the MAF for all three cell lines was greater than this value. All results from the screen are reported as MAF.

#### Targeted RNA-seq

RNA extraction of *in vitro* cells was performed using BioRad SingleShot Cell Lysis kit (Bio-Rad, California, USA, Cat #1725080) scaled down to 50 µl reactions per well of 96-well plate per manufacturer’s instructions. Tissue culture cells were lysed directly on the tissue culture plate. For RNA extraction of *in vivo* studies, tissues were harvested and placed in RNA Later (Invitrogen, Maryland, USA, Cat # AM7020) and stored at -80°C until RNA extraction. RNA extraction of tissues was performed using a Qiagen RNeasy Kit (Qiagen, Maryland, USA, Cat #74104) per manufacturer’s instruction with tissue homogenization in 600 µl of RLT buffer via dounce homogenizer followed by QIAshredder (Qiagen, Maryland, USA, Cat #79656). All RNA quality was assessed by Agilent Tapestation 2200 (Agilent, California, USA, Cat #G2964AA) and the Agilent RNA ScreenTape (Agilent, California, USA, Cat #5067-5576) with Agilent RNA ScreenTape Sample Buffer and Ladder (Agilent, California, USA, Cat #5067-5577, Cat #5067-5578) per manufacturer’s instruction. Reverse transcription to cDNA was performed using High-Capacity cDNA Reverse Transcription Kit (Applied Biosystems, USA, Cat #4368814) per the manufacturer’s instructions. cDNA was amplified using cDNA-specific primers with at least one primer (forward or reverse) covering an exon-exon boundary (See Table S6 for primer sequences). Primers were obtained from Integrated DNA Technologies (Iowa, USA). PCR was performed using Phusion Flash High-Fidelity PCR Master Mix (Thermo Fisher, USA, Cat #F548S) for 1 min at 98 °C, 30 cycles of 10 sec at 98 °C, 15 sec at 64 °C, 15 sec at 72 °C, and 5 min at 72 °C. Sequencing libraries were prepped from samples by addition of well barcodes using the same method described above for an additional 2-6 cycles. Libraries were pooled, cleaned up with AMPure beads (Beckman Coulter, California, USA, Cat #A63880) and sequenced on an Illumina Miseq using manufacturer’s instructions (150 cycle single read).

#### Mutant Allele Fraction (MAF) analysis

Mutant allele fraction was determined by processing fastq files using HISAT2 (version 2.0.5) and aligning to a pseudo reference genome consisting of only the mutant or wild-type amplicon sequences for targeted regions. The mutant allele fraction was determined by taking a ratio of the number of mutant transcripts to the total number of transcripts from the region in question. Initial data processing was performed in MSSQL and Excel.

#### Whole transcriptome RNA-seq

LS180 or NCI-H358 cells were run in biological duplicate, treated with DMSO or 5µM LY3023414. For RNA extraction, cells were pelleted, frozen in liquid nitrogen, and stored at -80°C until RNA extraction. RNA extraction was performed using a Qiagen AllPrep DNA/RNA Mini Kit (Qiagen, Maryland, USA, Cat# 80204) per manufacturer’s instruction with cell homogenization and lysis in RLT buffer with a QIAshredder (Qiagen, Maryland, USA, Cat# 79656). RNA quality control using Agilent Tapestation 2200 (Agilent, California, USA, Cat# G2964AA) and the Agilent RNA ScreenTape (Agilent, California, USA, Cat# 5067-5576) with Agilent RNA ScreenTape Sample Buffer and Ladder (Agilent, California, USA, Cat# 5067-5577, Cat# 5067-5578) per manufacturer’s instruction. Library prep using Illumina RNA library prep kit (Illumina, California, USA, Cat #RS-122-2001) and sequenced on an Illumina HiSeq 4000 150 cycle paired-end using manufacturer’s instructions.

#### RNA-seq analysis

Sequencing reads aligned to Hg38 using HISAT2 (version 2.0.5), RNA alignment metrics using CollectRnaSeqMetrics (Picard, version 2.20.2). Exon skipping was determined using IGV Viewer Sashimi Plots[84]. The average number of bases sequenced per sample and percent aligned in LS180 was 5.14e9 bases (range 5.07e9-5.20e9) and 77.8% (range 76.1%-78.7%) and for NCI-H358 was 5.58e9 bases (5.19e9-5.87e9) and 79.4% (range 77.2%-82.7%). Mean allele frequency was determined using VarScan 2[85] by generating the ratio of Read 1 (mutant) to read2 (wild type) transcripts. Mutations were only considered if they were heterozygous, and contained at least 5 reads at the somatic mutation or indel site in all 4 samples being compared (biological duplicates of treated and untreated).

#### Compound response curves

We performed a 6-point dose-response curve by treating cell pools for 14 hours with single compounds. Cell pools consisted of three isogenic knockout cell lines grouped by parental cell line with readout via targeted sequencing RNA (see *Targeted RNA-seq* for details). The effect of the compound was determined by calculating the MAF by comparing the abundance of the expected mutation and compared to the wild type within each isogenic pool (see *Mutant Allele Fraction (MAF) analysis* for details).

#### Immunohistochemistry (IHC)

IHC was performed on cell lines as previously described[86].

#### Western blots

Cells were lysed using RIPA buffer (Thermo Fisher, USA, Cat #89901) containing 1x protease inhibitor (Thermo Fisher, USA, Cat #4693159001) on ice for 30 minutes. Samples were then centrifuged at max speed for 3 minutes at 4°C in a QIA shredder (Qiagen, Maryland, USA, Cat #79654) before being transferred to a new sample collection tube. Protein was quantified using a BCA assay (Thermo Fisher, USA, Cat #23227) per the manufacture’s instructions. Gels were run by loading 50 µg of total protein per sample into 15 well polyacrylamide gels (Bio-Rad, California, USA, Cat #456-1086) and run for 30 minutes at 200V. Gels were then transferred using the manufacturer’s instructions (based on size) to nitrocellulose membrane using a Bio-Rad turbo transfer apparatus (Bio-Rad, USA, #170-4270).

Membranes were blocked for 1 hour with 3% milk-TBS-Tween before being incubated overnight in primary antibody (concentration dependent on antibody). Primary antibodies and concentrations can be found in the key resources table. Membranes were washed 4 times for 5 minutes each with TBS-Tween. Secondary antibody was applied at 1:2500 using either α-rabbit (Jackson ImmunoResearch, Pennsylvania, USA, Cat #111-035-006) or α-mouse (Jackson ImmunoResearch, Pennsylvania, USA, Cat #115-035-006). Membranes were imaged using Pierce™ ECL Western Blotting Substrate (Thermo Fisher, USA, Cat #32106) following the manufacturer’s instructions on a Bio-Rad Chemidoc (Bio-Rad, California, USA). Phospho-UPF1 westerns were performed as detailed above with the following exceptions: secondary antibodies were used at 1:10,000 either α-rabbit (Licor Biosciences, USA, Cat #926-32213) or α-goat (Licor Biosciences, USA, Cat #926-68074). Membranes were imaged using an Odyssey CLx (Licor Biosciences, USA) following the manufacturer’s instructions.

#### siRNA

We performed knockdown of kinase proteins using siRNA. LS180 and NCI-H358 cells were plated in 96-well plates (Costar, USA, Cat #3595) at 5,000 cells per well. Cells were transfected with DharmaFECT1 transfection reagent (Horizon, USA, Cat #T-2001-02) and either 50 nM control or kinase-specific pooled siRNA. siRNA used in this experiment can be found in the key resources table. Knockdown efficacy was confirmed using qPCR (see *Quantitative real-time PCR (qPCR)* for details). NMD target gene mutation transcript levels were determined using next-generation sequencing and MAF (see *Targeted RNA-seq* for details).

#### Bispecific scFv construction

We utilized a bispecific antibody against CD-3 and TP53 wild-type peptide ‘TYSPALNKMF’ (residues 125-134) presented in a HLA-A24 MHC-I molecule scFv. This bispecific antibody was identified by panning a phage display library. The scFv-bearing phage library was constructed similarly as described in detail previously with some modifications[87]. Briefly, oligonucleotides were synthesized by GeneArt (Thermo Fisher, USA) using trinucleotide mutagenesis (TRIM) technology to diversify complementarity-determining region (CDR)-L2, CDR-L3, CDR-H1, CDR-H2, and CDR-H3. A FLAG (DYKDDDDK) epitope tag was placed immediately downstream of the scFv, which was followed in frame by the full-length M13 pIII coat protein sequence. The total number unique clones obtained was determined to be 3.6 x 10^10^. Panning details can be found in the reference section[86, 87].

#### Kinase target assay

Kinase kinativ experiments were performed by ActivX Biosciences (San Diego, USA)[54].

#### Quantitative presentation of HLA-bound neoantigens via HPLC-Mass Spectrometry

Identification and quantitation of HLA-presented neoantigens was performed as previously described by Complete Omics Inc.[57]. Briefly, cells were treated *in vitro* for 24 hours with either DMSO or 5 µM KVS0001. Cells were cross-linked and immunoprecipitated with pan-HLA antibodies to obtain cell surface MHC-presented peptides. Mass spectrometry was performed in the presence of heavy labeled peptide to serve as an internal (loading) control to quantify the presence of the presented neoantigen. The presented peptides were identified in a preliminary MS screen of 187 candidate peptides predicted by using the union of NetMHC and Predictor of Immunogenic Epitopes (PRIME) predictions[58, 59].

#### CD3-TP53scFv bispecific co-culture

Co-culture of bispecific antibody was performed using volunteer human donor T-Cells. T-cell enrichment and activation was performed as previously described [88]. Briefly, PBMCs are incubated with OKT3 antibody (Biolegend) for 3 days. T-cells were then expanded in RPMI-1640 with 10% FBS and 1% penicillin-streptomycin, recombinant IL-2 (Proleukin, Prometheus Laboratories) and IL-7 (BioLegend) for at least 15 days before use. NCI-H716 cells were labeled with HLA-A24 using retrovirus as previously described[86]. Briefly, the MSCV retroviral expression system (Clontech, USA, Cat #634401) was used to overexpress HLA-A*24-T2A-GFP in target cells. Expression was confirmed by flow cytometry. NCI-H2228 cells were not labeled as they express this specific HLA endogenously. For NCI-H716_A24 and NCI-H2228, 40,000 cells were plated in 96-well plates, and co-incubated with 40,000 activated T-cells, and 12.5 pg/mL of TP53-CD3 bispecific antibody. Cells were then dosed with 1.25µM of KVS0001, SMG1 specific inhibitor or 200 mg/mL of 5-fluoruracil (Sigma Aldrich, USA, Cat #F6627) for 24 hours. Readout of interferon (IFN-γ) was performed by ELISA following the manufacturer’s instructions (R&D Systems, USA, Cat #SIF50C). Cell killing was assayed by luciferase levels following the manufacturer’s instructions (Promega, USA, Cat #E1501) and read out on a Synergy H1 Microplate reader (BioTek, USA). NCI-H716_A24 cells were labeled with luciferase via lentiviral transduction following the manufacturer’s recommendations (OriGene, USA, Cat #PS100071).

#### Animal protocols

Animal research was approved and overseen by Johns Hopkins University Institutional Animal Care and Use Committee (ACUC) approved research protocol M018M79. Mice are housed in individually ventilated caging (Allentown, New Jersey, USA) at a maximum 5 animals per cage. Cages are changed every 14 days. Enrichment is provided through paper bedding, paper hut, and some food placed in the bottom of the cage. Facility is maintained between 70 and 72 degrees Fahrenheit on a 12-hour light-dark cycle. Mice standard diet is ad lib Teklad Global 18% Protein Extruded Rodent Diet, autoclaved (Envigo, Huntingdon, UK) and acidified water via sipper tube.

#### In Vivo tumor models

Cells for tumor inoculation in each mouse were grown to 70-80% confluency *in vitro*. Cells were harvested with trypsin (Gibco, California, USA, Cat #25300054) and suspended in either PBS for LS180 cells or 50% PBS, 50% Matrigel Phenol Red Free Standard Formulation (Corning, New York, USA, Cat #356237) for NCI-H358 cells. For both cell lines 1e6 cells were placed subcutaneously and grown to approximately 200 mm^3^. Animals were randomized into treatment groups prior to treatment by tumor size. All tumor volumes were measured via caliper twice weekly. Mice with tumors <150 mm^3^ at 11 days post inoculation were excluded from experiments.

#### Human xenograft experiments

6 to 8 weeks old *Mus musculus* Hsd:Athymic Nude-Foxn1^nu^ mice (referred to as nude mice) were purchased from Harlan Laboratories (Indiana, USA). Only female mice were used as gender was not considered to be a significant confounder in the experiment. LS180 cells were inoculated at 1.0 x 10^6^ cells per mouse in the left mouse flank or NCI-H358 cells were inoculated at 7.5 x 10^5^ cells per mouse in the right flank. For single-dose experiments, mice were orally dosed via gavage at 0, 40, or 60 mg/kg of LY3023414 (Selleckchem, Texas, USA, Cat #S8322) in 1% Methylcellulose (Sigma Aldrich, Missouri, USA, Cat #M6385). KVS0001 was dosed at 30 mg/mL intraperitoneal (IP) in 0.5% Dimethyl sulfoxide (DMSO)(Sigma Aldrich, USA, Cat #C6295), 10% cremaphor (Sigma Aldrich, USA, Cat #C5135) and 2% glycerol (Sigma Aldrich, USA, Cat #G5516). Mice were given physical exams prior to euthanasia at designated endpoints according to the approved research protocol. Tumors, spleen, blood, and lungs were harvested post-euthanasia for further analysis. Tumors were harvested at indicated times and stored in RNA Later (Invitrogen, Maryland, USA, Cat # AM7020) at -80 for further analysis.

#### Syngeneic mouse tumor experiments

6 to 8 weeks old immune compromised *Mus musculus* BALB/cAnNCrl (referred to as BALB/c) and C57BL/6NCrl (referred to as C57BL/6N) mice were purchased from Charles River GmbH (Germany). Only female mice were used as gender was not considered to be a significant confounder in the experiment. Murine cancer cell lines were placed in the left mammary fat pad of female mice at the following densities: LLC 1.0 x 10^6^, RENCA 1.0 x 10^6^, and B16-F10 0.2 x 10^6^. Mice were randomized around day 7 post-implantation when tumor sizes had reached approximately 30-40 mm^3^. Mice were dosed with 30 mg/kg KVS0001 or vehicle control via IP injection alternating the right and left side daily for 28 days (Figure 5C). Mice were given physical exams before euthanasia at designated endpoints according to the approved research protocol.

#### In Vivo TP53 bispecific experiments

For the NCI-H716 *in vivo* bispecific experiments 2.5 x 10^6^ NCI-H716_A24 expressing cells were placed orthotopically by IP injection into NOD.Cg-Prkdcscid Il2rgtm1Wjl/SzJ (referred to as NSG) mice (Jackson Laboratories, USA). On day 2 mice were randomized to ensure a balanced tumor burden in all groups. Luminescence was measured by injecting mice with 150 µL of RediJect D-Luciferin Ultra Bioluminescent Substrate (PerkinElmer, USA, Cat #770505), and anesthetized using isoflurane in an induction chamber for 5 minutes. Readouts and analysis were performed on an IVIS Spectrum imaging system and Living Image software (Perkin Elmer, USA).

#### Data reporting

No statistical methods were used to predetermine the sample size. Statistical analysis was performed using R and Excel. All animal experiments were randomized being sure to alter the first cage dosed and location of cages with a row of the rack. Randomization was performed by animal tumor size, unless otherwise indicated. The investigators responsible for weight and tumor measurements were blinded to the allocation, treatment, and outcome assessment of experiments. The investigators processing mouse tissue processing were blinded to the allocation, treatment, and outcome assessment of experiments. The investigator harvesting the tissues was aware of the allocation, treatment, and outcome assessment of experiments.

#### Statistical Testing

Chi-Squared testing was performed using counts of cancer cell line mutations predicted to undergo NMD and RNA recovered and prop.test() in R. Mann-Whitney testing performed with wilcox.test() in R. Quantiles calculated utilizing z scores with quantile() in R. Student T-Test and one-way ANOVA with Dunnett’s test and Student’s t-test were performed in R and Excel.

---

**Figure S1:**
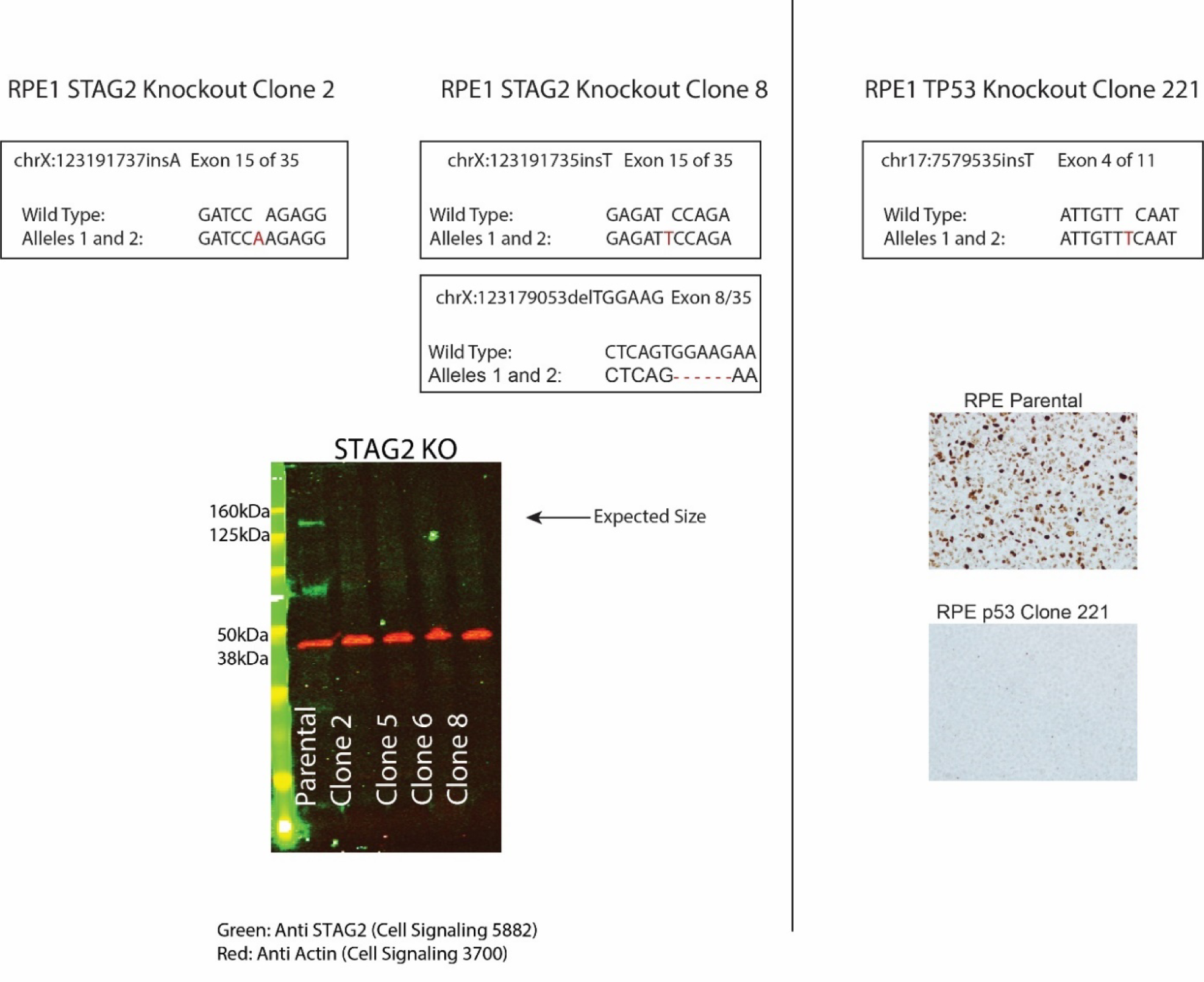
Next generation sequencing results depicting genomic mutations at the CRISPR target area in *STAG2* (top left) and *TP53* (top right) isogenic cell line clones in the RPE1 cell line which were used in subsequent experiments. Western blot showing STAG2 protein loss in four independent clones of which only clones 2 and 8 were used in subsequent experiments (bottom left). Note that STAG2 clone 8 has two independent biallelic indel events. The mutation in exon 15 was used for scoring relative transcript levels in the screen and for subsequent work. IHC results confirming p53 protein loss in RPE TP53 221 (bottom right).

**Figure S2:**
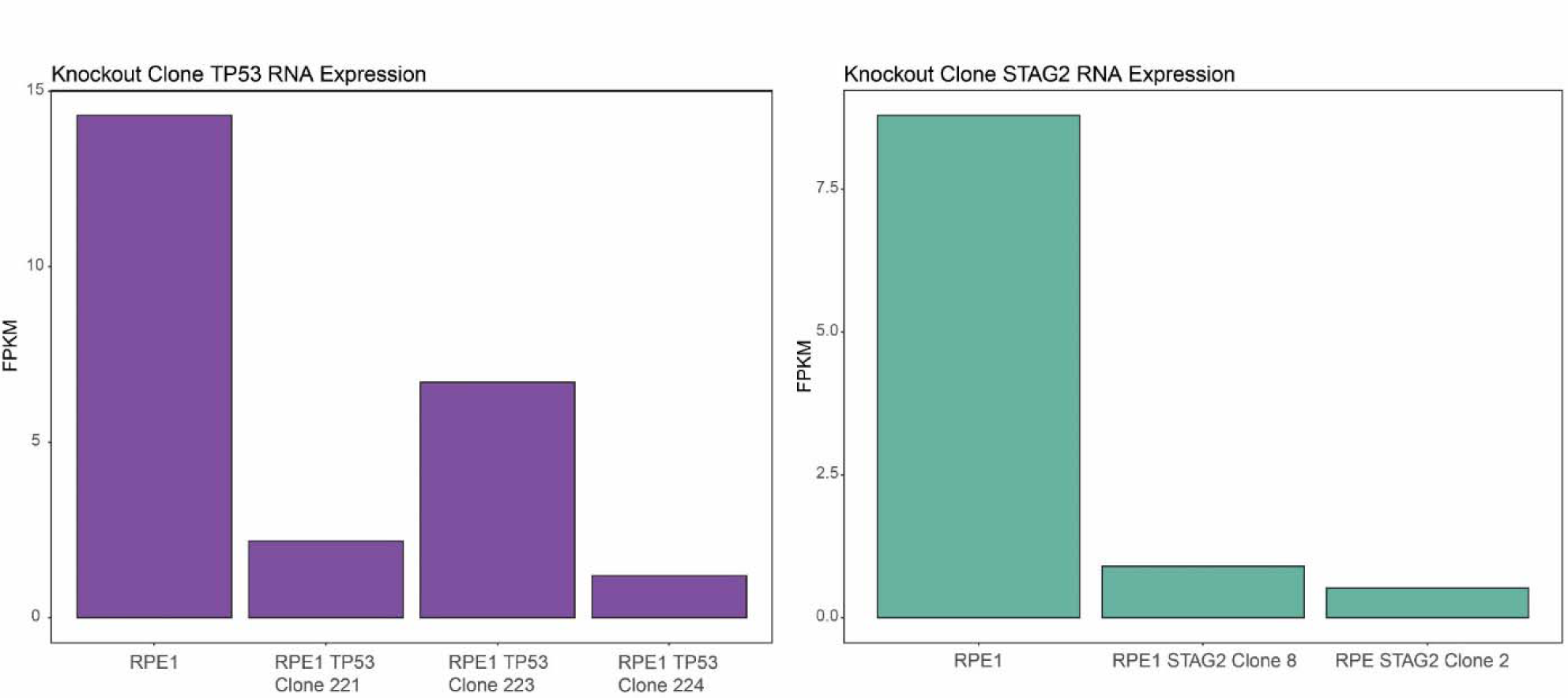
Fold change in RNA expression levels from whole transcriptome RNA-sequencing data for *STAG2* and *TP53* knockout clones in the RPE1 cell line background. RPE1 TP53 clone 221, RPE1 STAG2 clone 2 and RPE1 STAG2 clone 8 were used in the HTS screen. RPE1 TP53 clone 223 and RPE1 TP53 clone 224 were used in Figure S6. Note clone 223 has a 9bp in-frame deletion in one allele and an out of frame deletion on the other allele, presumably accounting for the higher level of expression.

**Figure S3:**
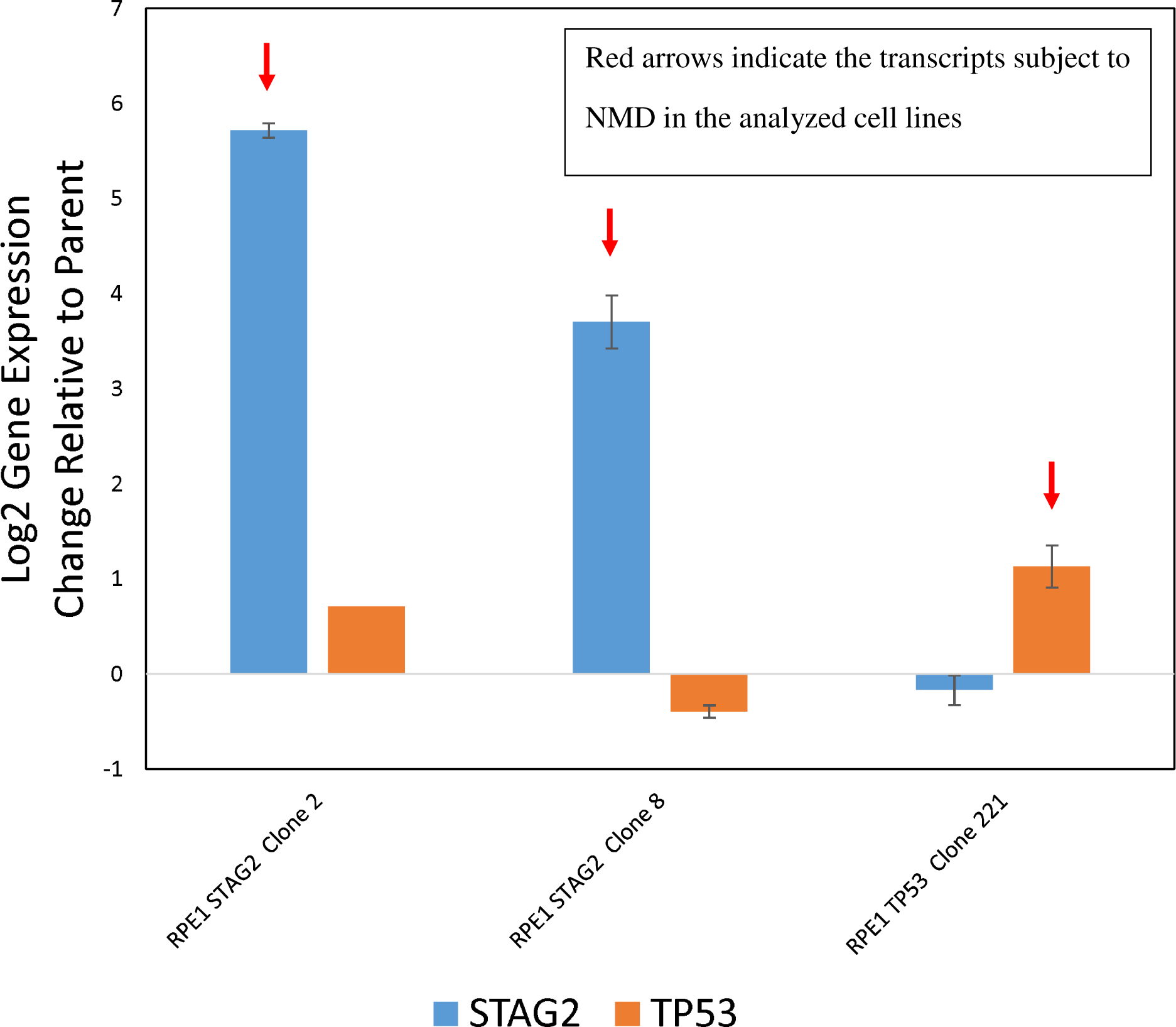
RNA transcript level changes based on qPCR in *STAG2* and *TP53* knockout clones treated with the known NMD inhibitor emetine at 12 mg/mL. Error bars show standard deviation of three biological replicates. All changes are statistically significant by student t-test (p<0.05).

**Figure S4:**
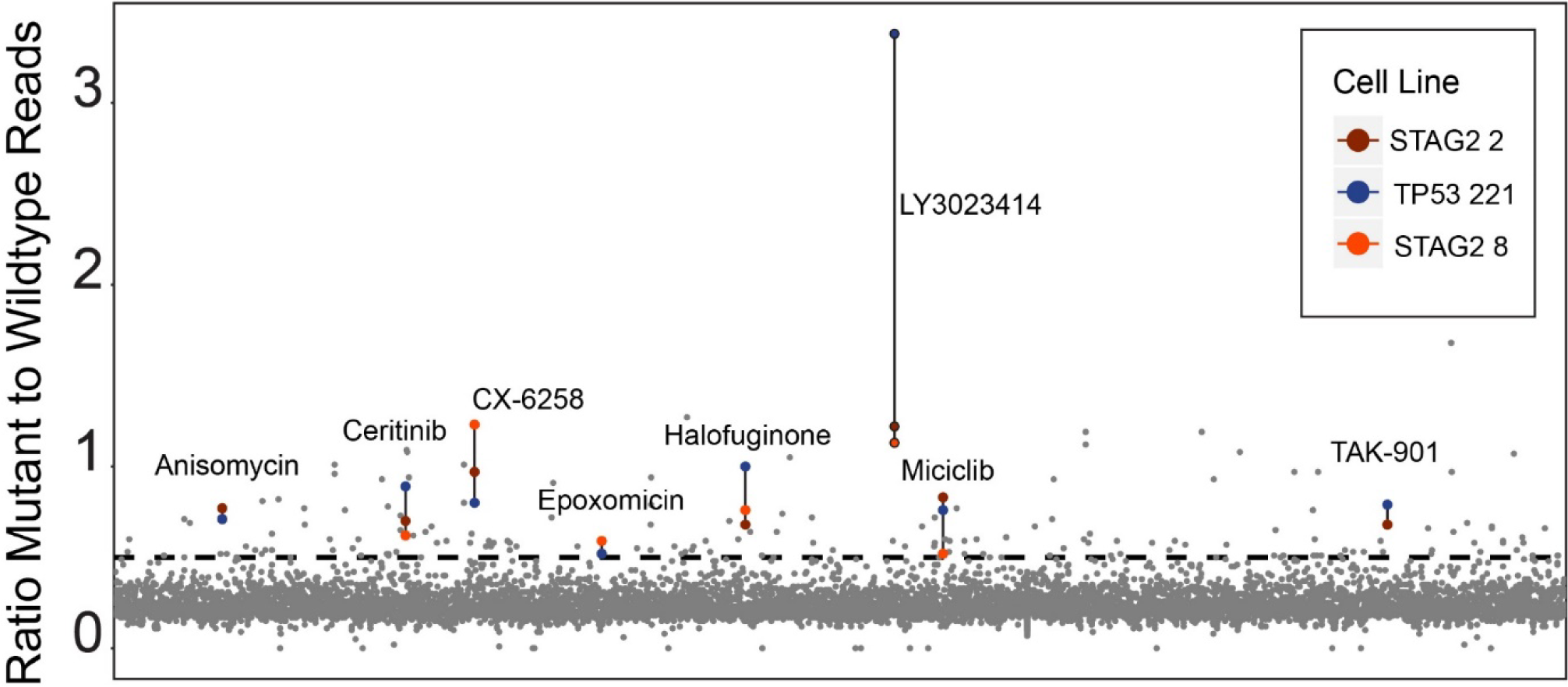
Primary screen results from high throughput assay. The x-axis represents all 2,658 compounds screened at 10µM, y-axis shows ratio of mutant to wild type reads for each of the three isogenic cell lines. Higher values indicate more mutant RNA reads, representing inhibition of NMD. The dotted line at 0.46 is the cutoff for a hit to be called (5 standard deviations above DMSO treated samples). Colored data points demarcate the eight hit compounds in the three screened cell lines.

**Figure S5:**
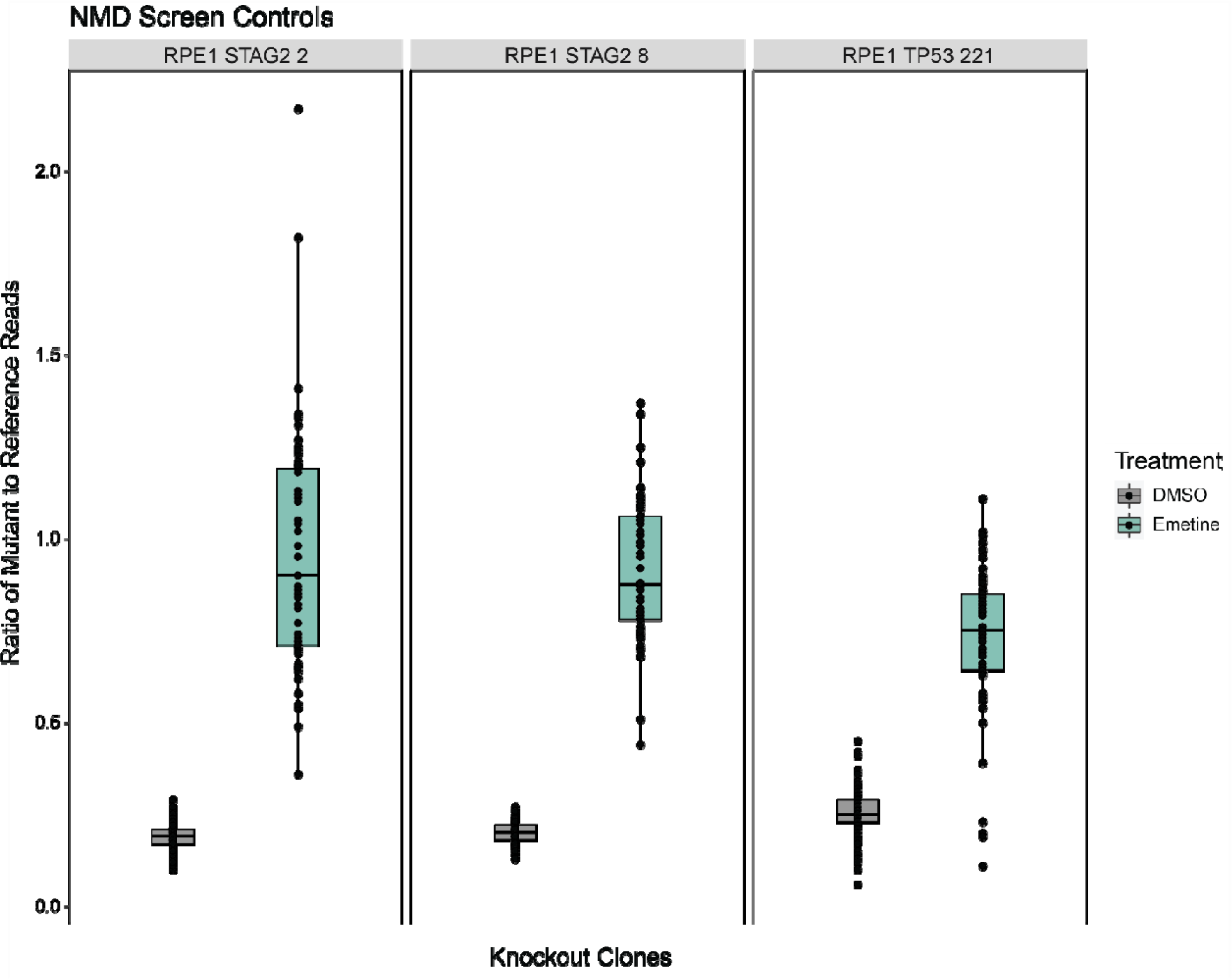
Emetine and DMSO control sample data from the HTS for each of the three clones used as measured by deep-targeted RNA sequencing. 6 DMSO and 2 emetine (12mg/mL) samples were included in each dosing plate for a total of 198 DMSO and 66 emetine measurements in each boxplot.

**Figure S6:**
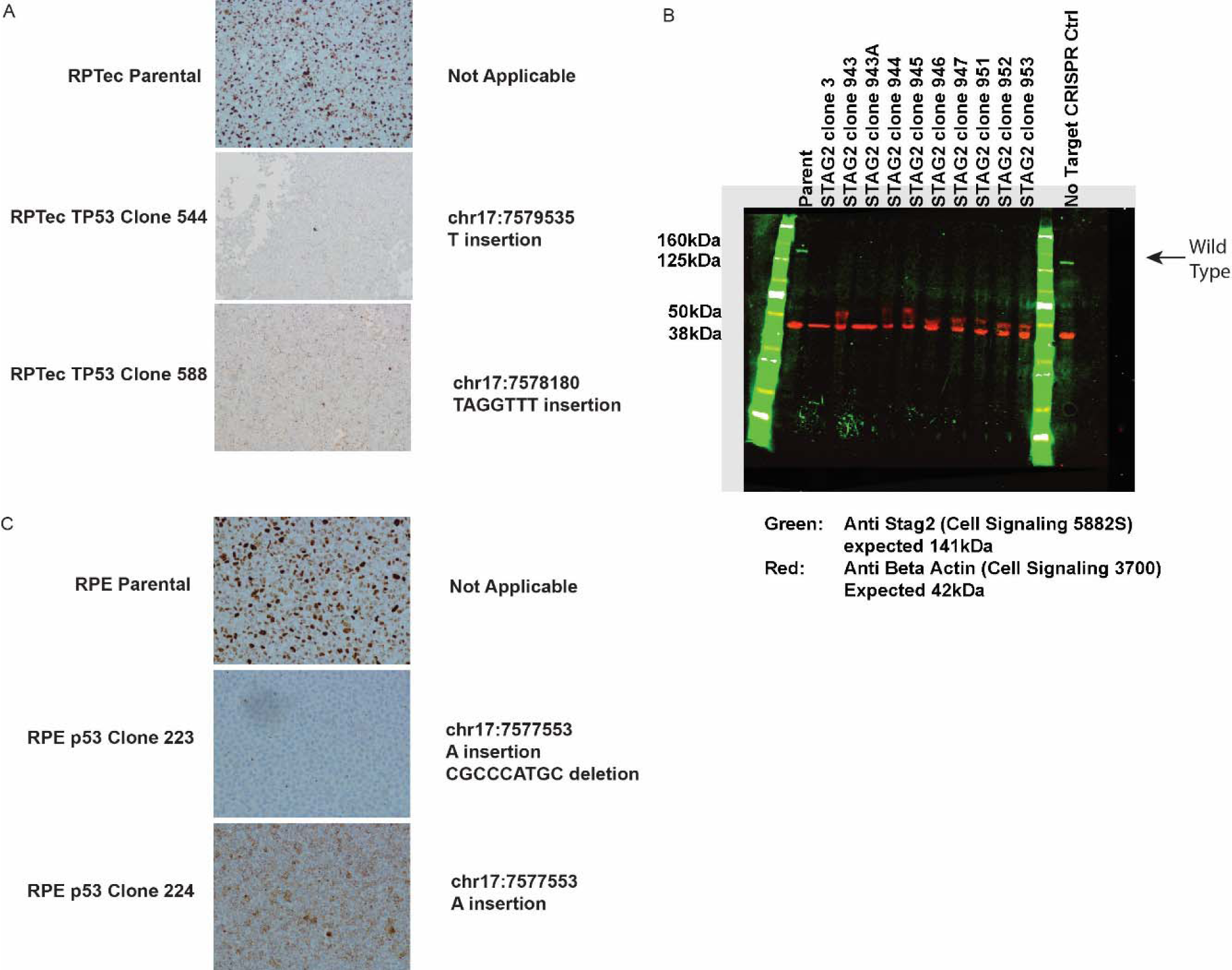
A) IHC staining of TP53 in RPTec *TP53* knockout clones. **B)** Western blot against STAG2 on RPTec *STAG2* knockout clones demonstrating successful knockout at the protein level of all ten clones. Out of frame indels were confirmed in all clones by NGS (data not shown). The arrow indicates the expected size for full-length STAG2 protein. **C)** IHC staining of TP53 protein on RPE1 *TP53* knockout clones. Clone 223 contains the same truncating mutation found in clone 224 on allele 1 and has a 9 bp in-frame deletion that preserves some full length TP53α and TP53β isoform expression on allele 2 (See Figure 1E).

**Figure S7:**
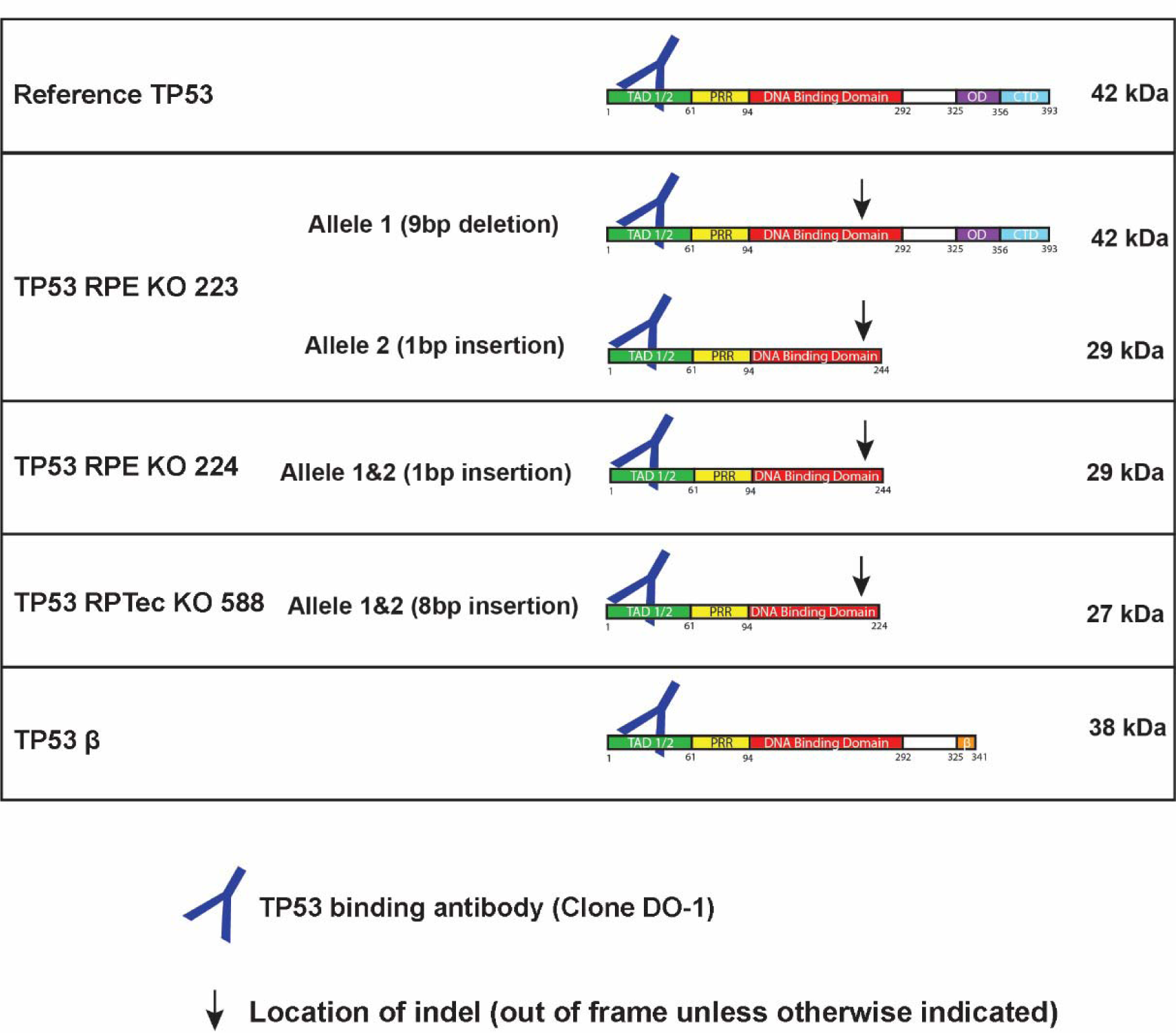
Protein schematic cartoons showing indel mutation site and expected size of various *TP53* knockout clones used in this study. Note, RPE TP53 223 has an in-frame deletion in the DNA binding domain.

**Figure S8:**
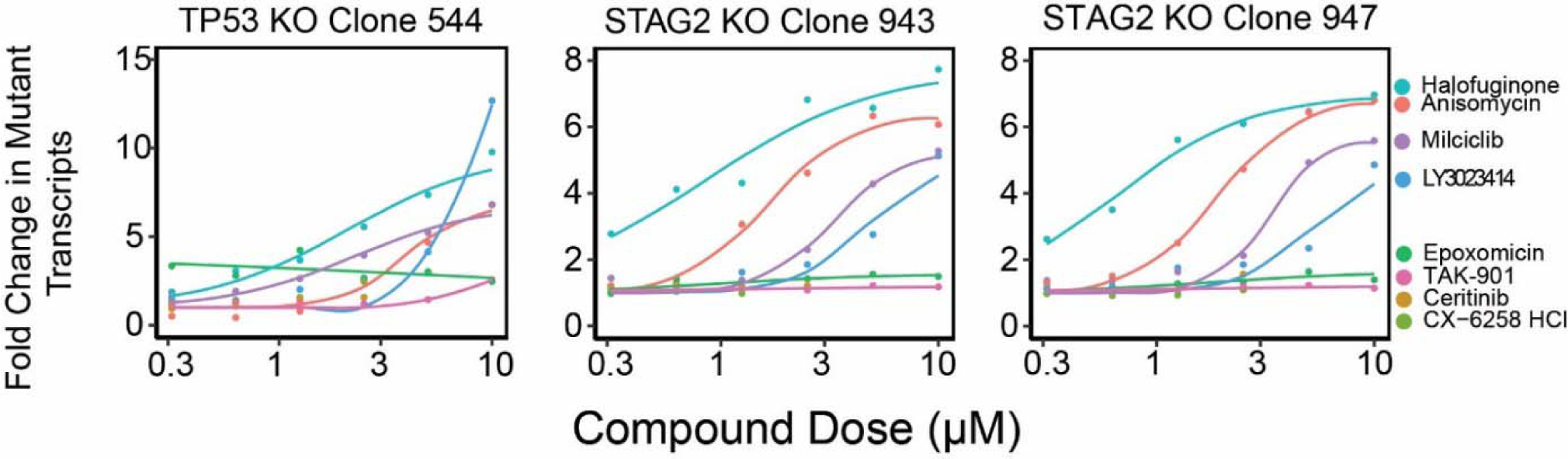
Fold change in mutant RNA transcription levels for STAG2 and TP53 in three knockout cell lines from the RPtec background containing out of frame indels targeted by NMD. Each line was treated with each of the eight hit compounds from the screen. The 10µM dose is also shown in figure 1C.

**Figure S9:**
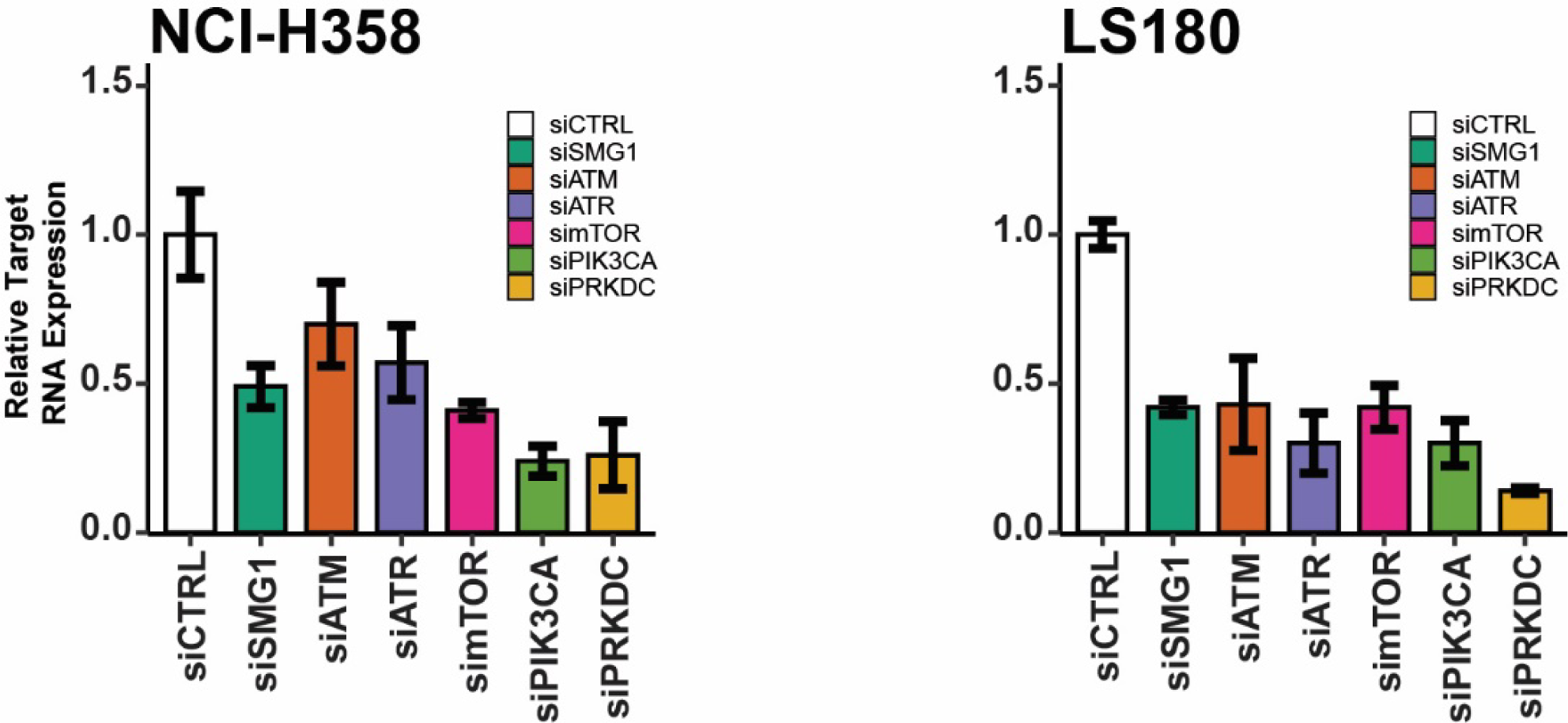
RNA expression levels of kinases post siRNA targeting in NCI-H358 and LS180 cells by qPCR. Cells are treated with siRNAs targeting genes known to be inhibited by LY3023414. Error bars represent 95% confidence intervals.

**Figure S10:**
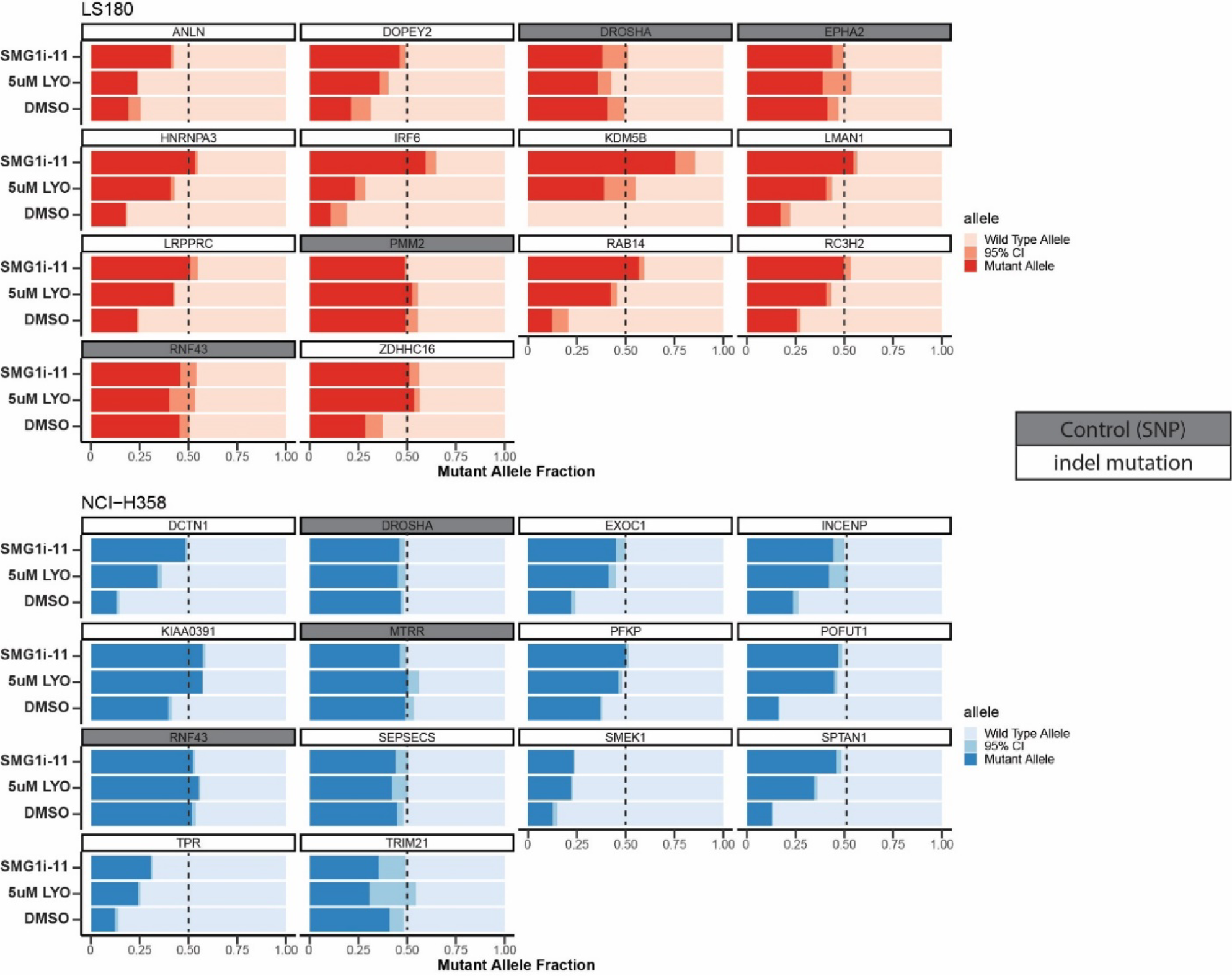
Slider plots showing mutant allele fraction relative to total reads measured by deep-targeted RNA-sequencing for *in vitro* treated LS180 (top in red) or NCI-H358 (bottom in blue) cells with 5µM LY3023414 (labeled LYO) or a previously described SMG1 inhibitor SMG1i-11 at 1µM. Gene names are shown in the boxes above each slider plot, genes highlighted in gray are common SNPs and serve as a negative control (not expected to change). In the case of the control SNPs, the mutant allele refers to the non-reference genome allele. *TRIM21* did not show a large change in expression with either small molecule, while *ANLN* did not show a difference with LY3023414 but did respond to SMG1i-11. The remaining genes responded to NMD inhibition by both LY3023414 and SMG1i-11.

**Figure S11:**
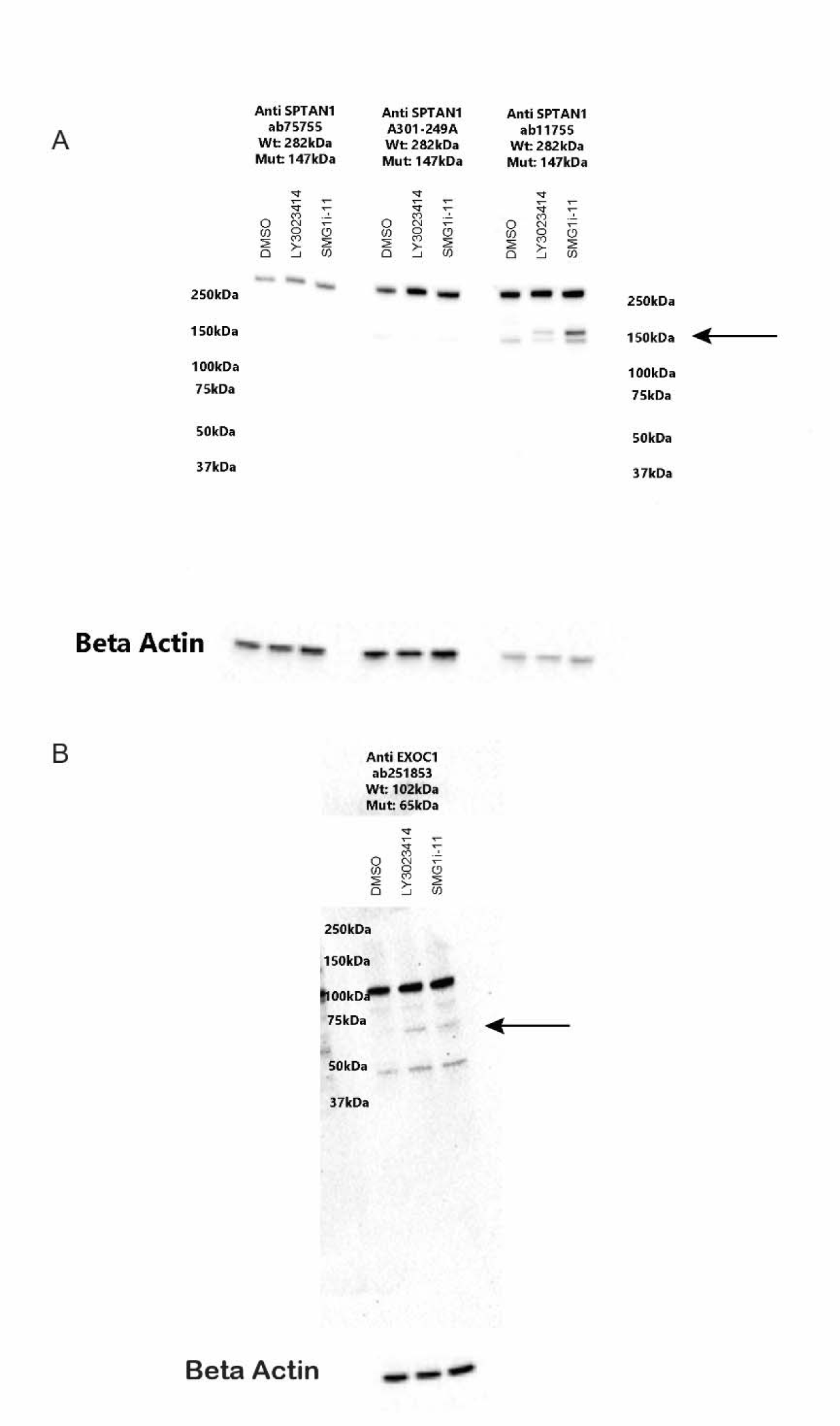
A) Western blot showing SPTAN1 expression after treatment with DMSO, 5µM LY3023414, or 1µM *SMG1* inhibitor SMG1i-11 (lanes 1,2,3 respectively for each antibody). The arrow indicates the expected size of the mutant (NMD targeted) protein. Antibody ab75755 (Abcam) binds C-terminal to the out of frame indel and does not show mutant protein as expected. Antibody A301-249 (Bethyl) is polyclonal and also did not bind mutant protein. Antibody ab11755 (Abcam) is located N-terminal to the indel and does display mutant protein expression. **B)** Western blot showing EXOC1 expression after treatment with DMSO, 5µM LY3023414, or 1µM SMG1i-11 (lanes 1,2,3 respectively). The arrow indicates the expected size of the mutant (NMD targeted) protein.

**Figure S12:**
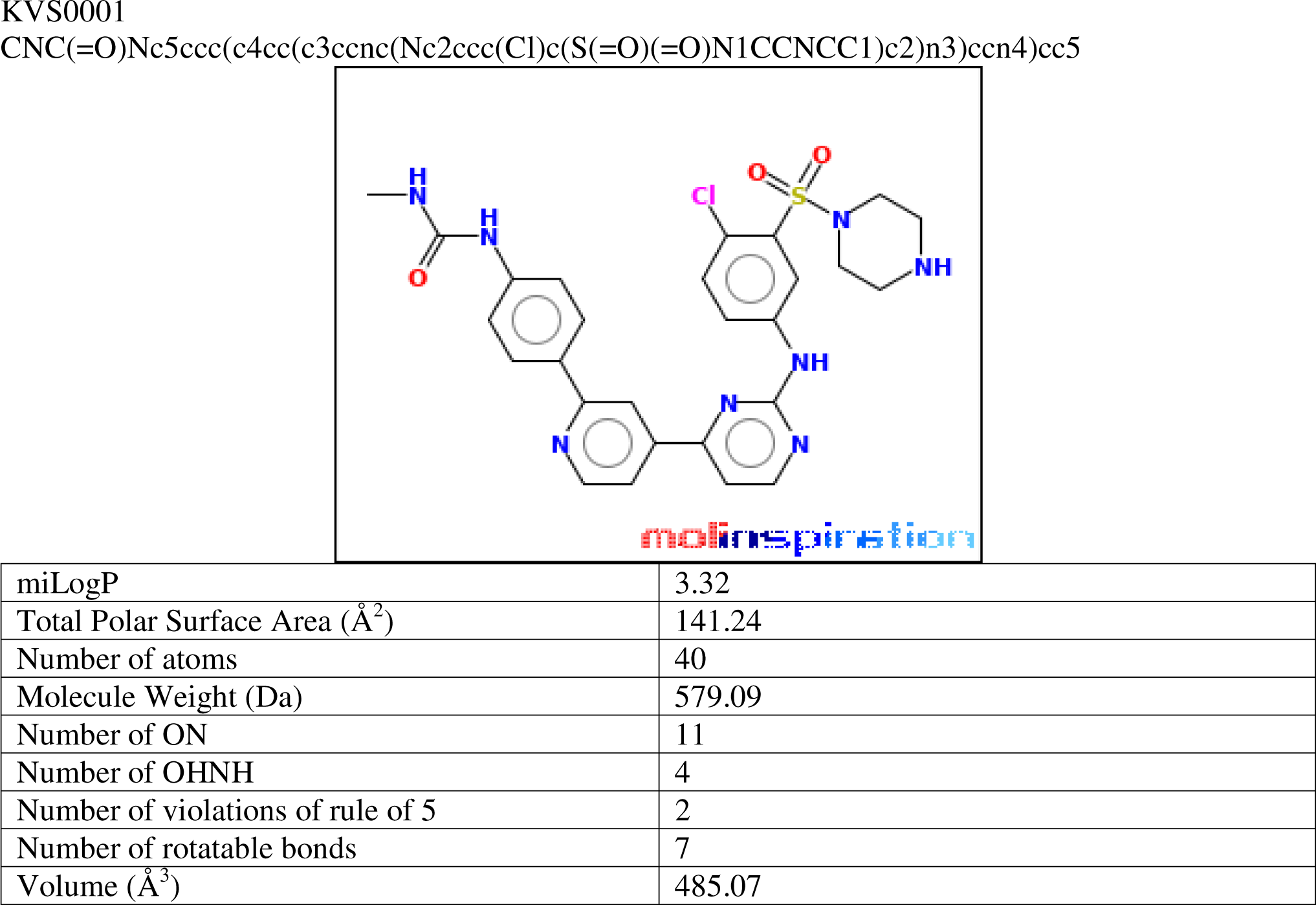
Biophysical properties for novel SMG1 inhibitor KVS0001.

**Figure S13:**
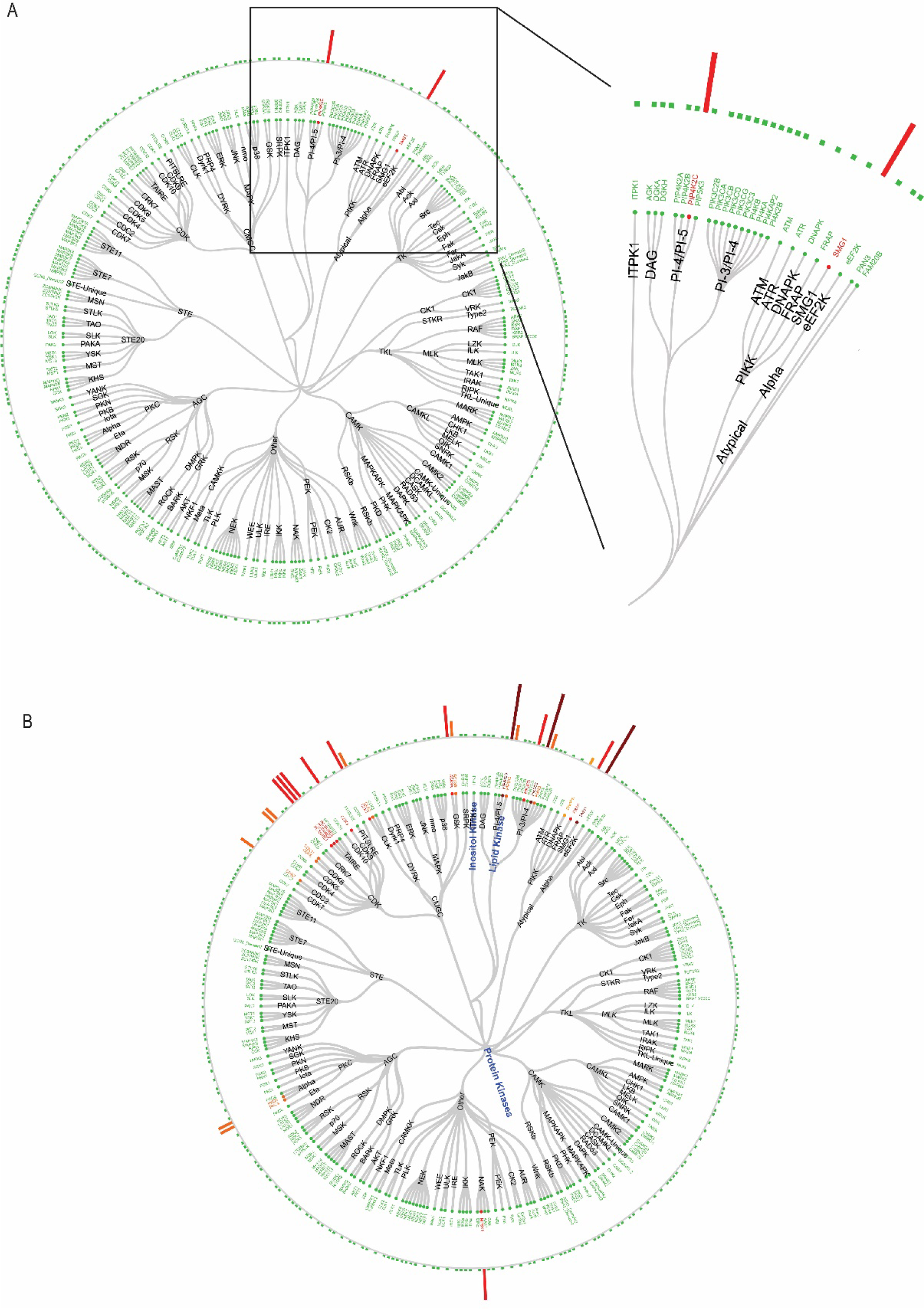
A) Kinativ™ Assay results for KVS0001 at 100 nM and B) 1µM run with biological replicates showing KVS0001 specificity against the known kinome. Results are based on the average between two unique peptides for each kinase.

**Figure S14:**
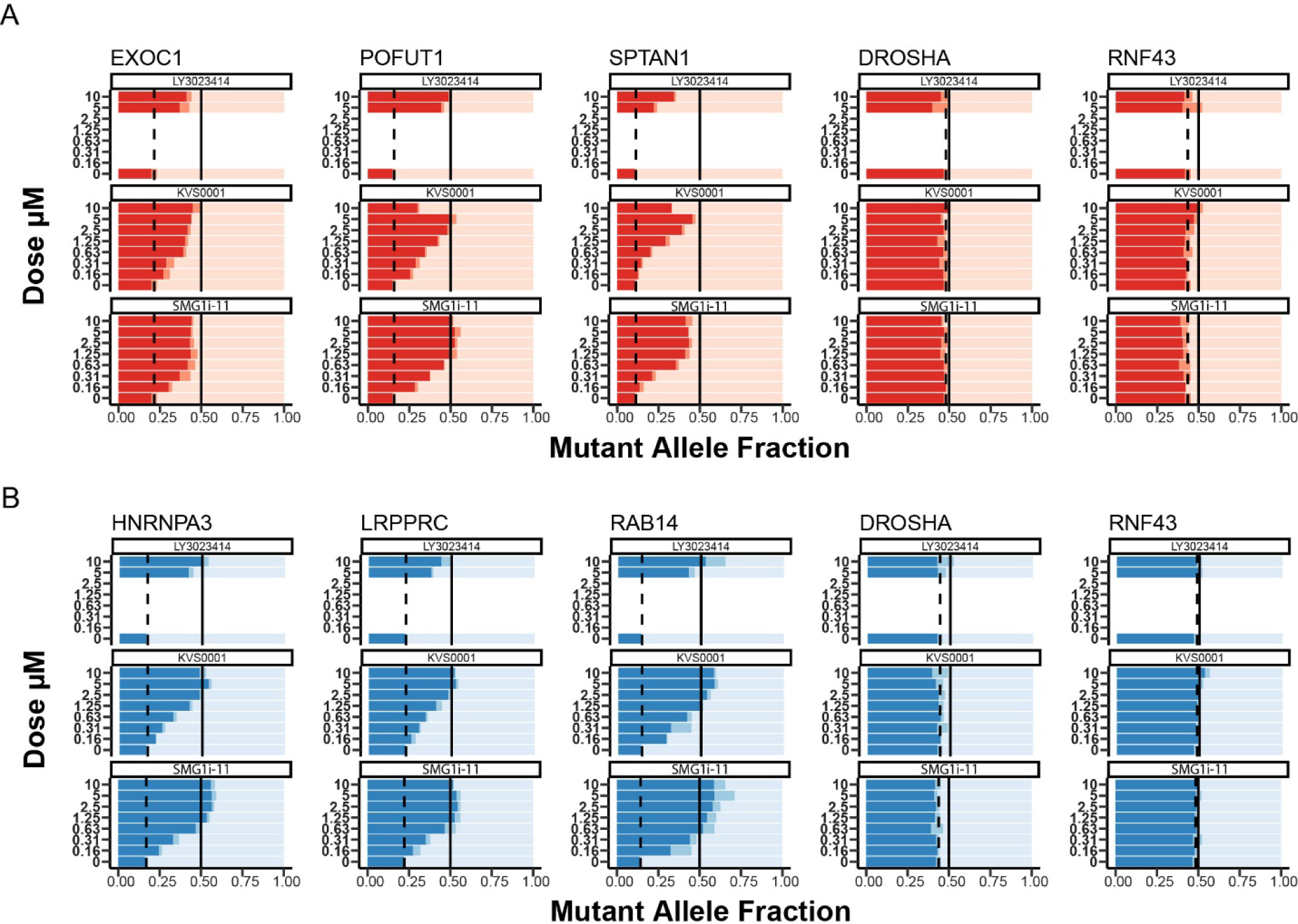
**A)** Slider plots showing mutant allele fraction measured by deep-targeted RNA-sequencing for genes from NCI-H358 (top in red) and **B)** LS180 (bottom in blue) treated in a dose response *in vitro* with novel NMD inhibitor KVS0001. *DROSHA* and *RNF43* are common heterozygous SNPs and serve as a negative control. In the case of the control SNPs, the mutant allele refers to the non-reference genome allele. Only the highest two concentrations were tested on LY3023414 which served as a positive control in this experiment. The dotted line indicates the mutant expression with DMSO treatment and the solid line is a reference for equal expression of both the wild type and mutant alleles.

**Figure S15:**
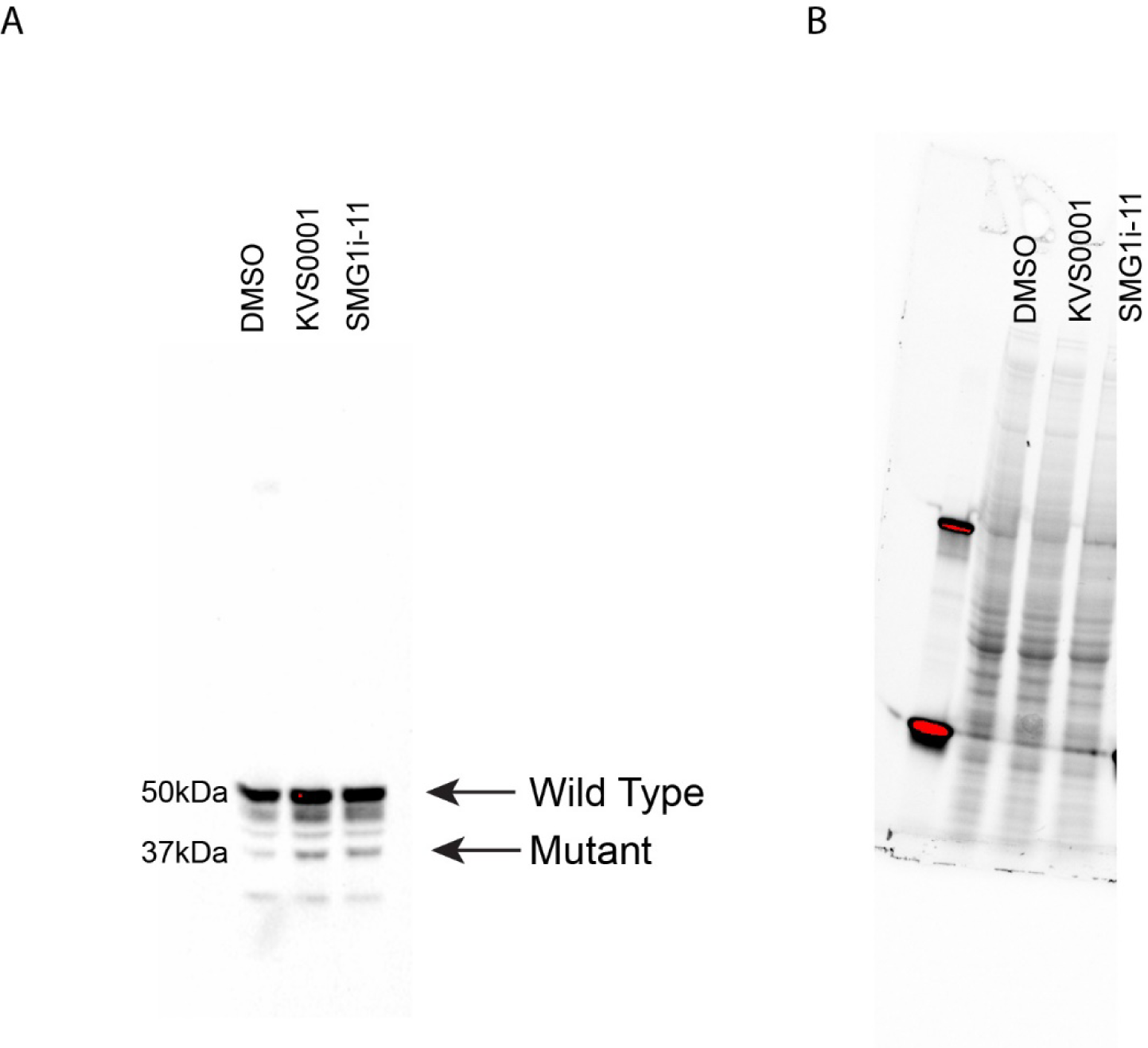
**A)** Western blot showing expression of LMAN1 in LS180 cells treated with DMSO (lane 1), KVS0001 at 5µM (lane 2) or SMG1i-11 at 1µM (lane 3). Arrows indicate expected size of wild type and mutant LMAN1 protein. **B)** Stain free loading control image for gel.

**Figure S16:**
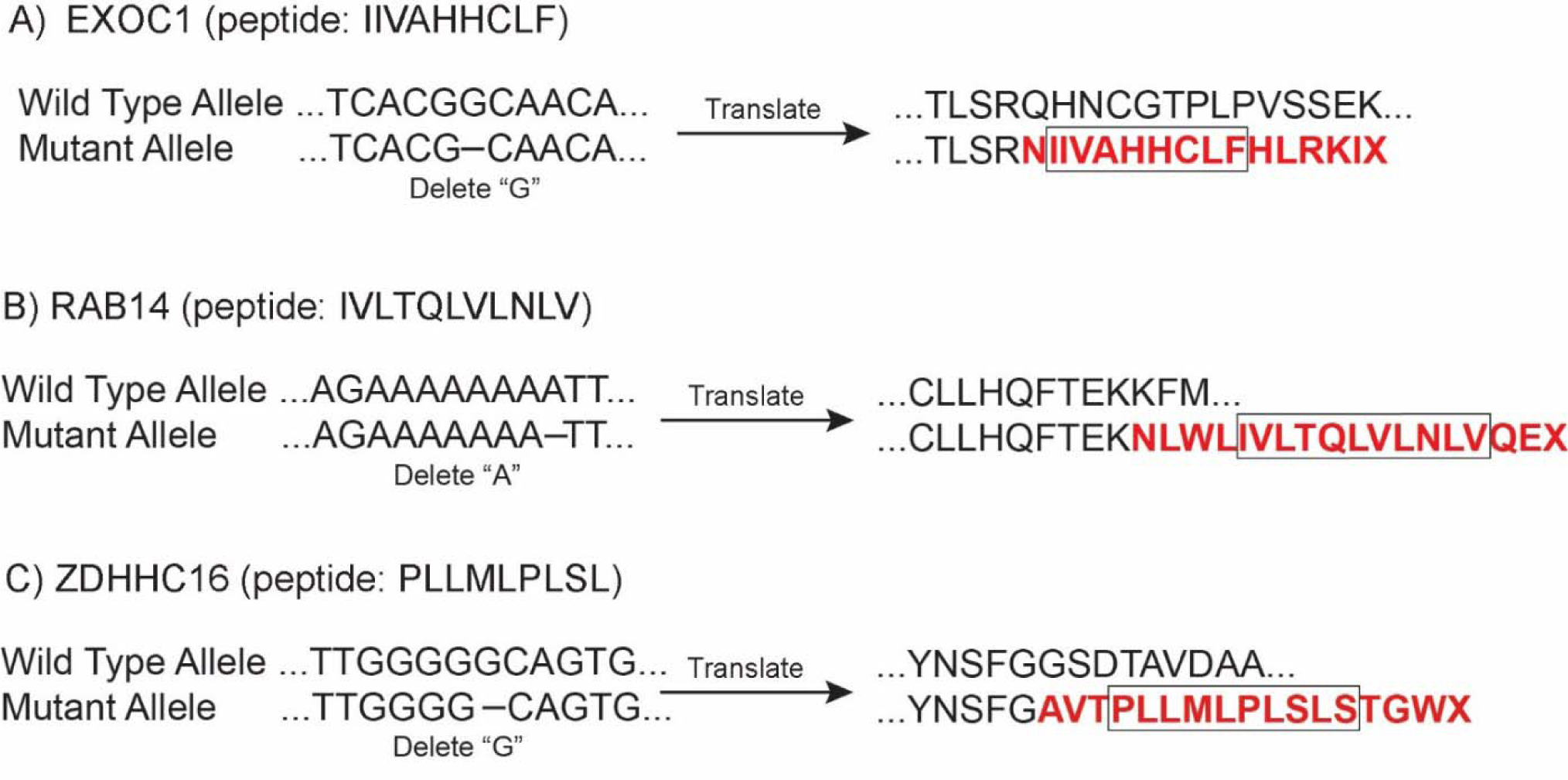
DNA and protein sequences for the wild type and mutant alleles of **A)** *EXOC1* **B)** *RAB14* and **C)** *ZDHHC16* genes. The mutant protein sequences caused by the out of frame indel are highlighted in red and the boxes indicate the peptides presented on the cell surface and identified by mass spectrometry in cells treated with 5µM of KVS0001 (see Figures 4A and 4B).

**Figure S17:**
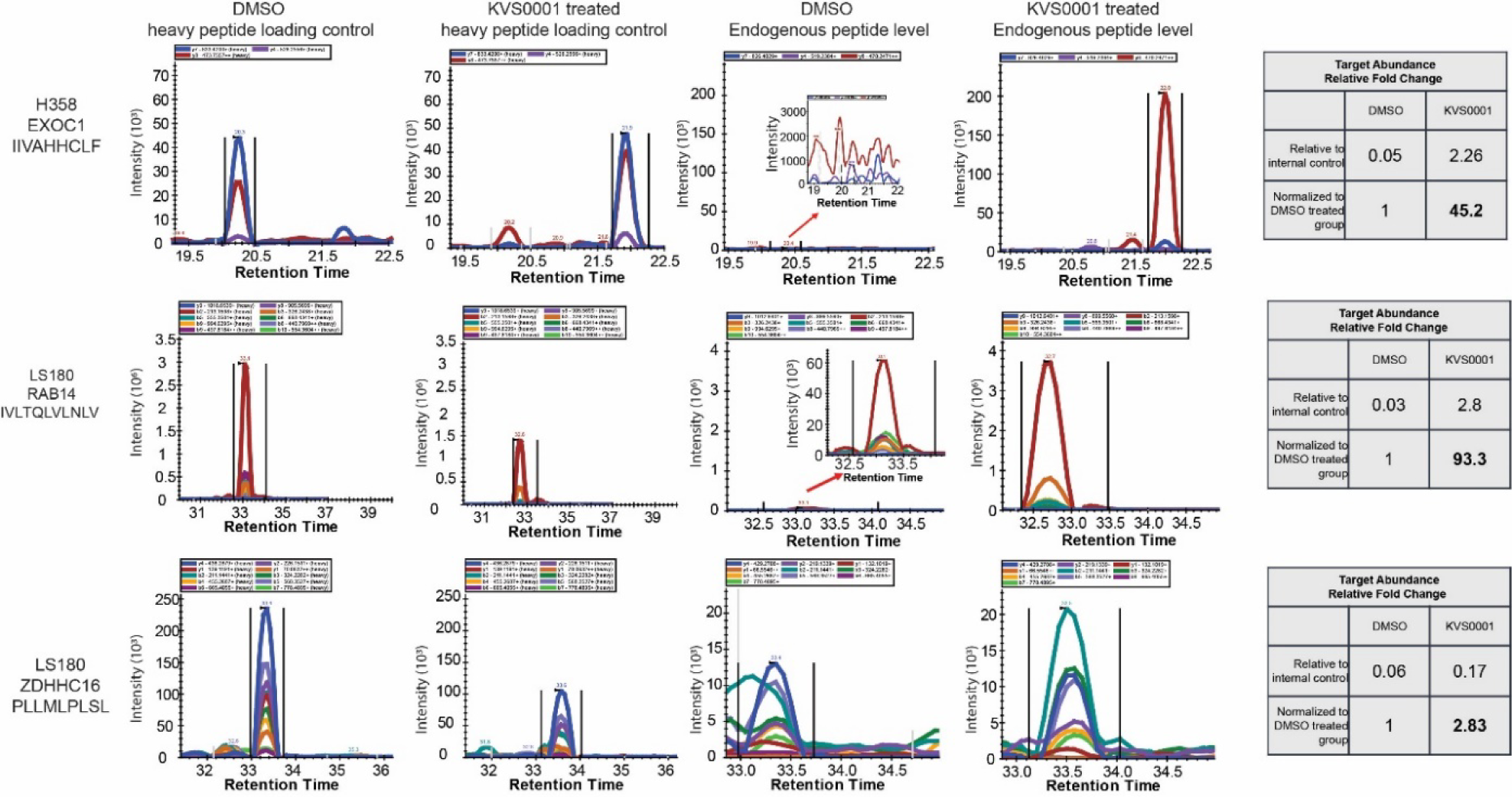
Heavy peptide loading controls and endogenous (light) peptide presentation of genes in LS180 and NCI-H358 treated with DMSO or 5µM KVS0001. Data is from quantitative HPLC-Mass Spectrometry. Note y-axis scale changes between samples. Tables on the right show relative increase in peptide presentation with KVS0001.

**Figure S18:**
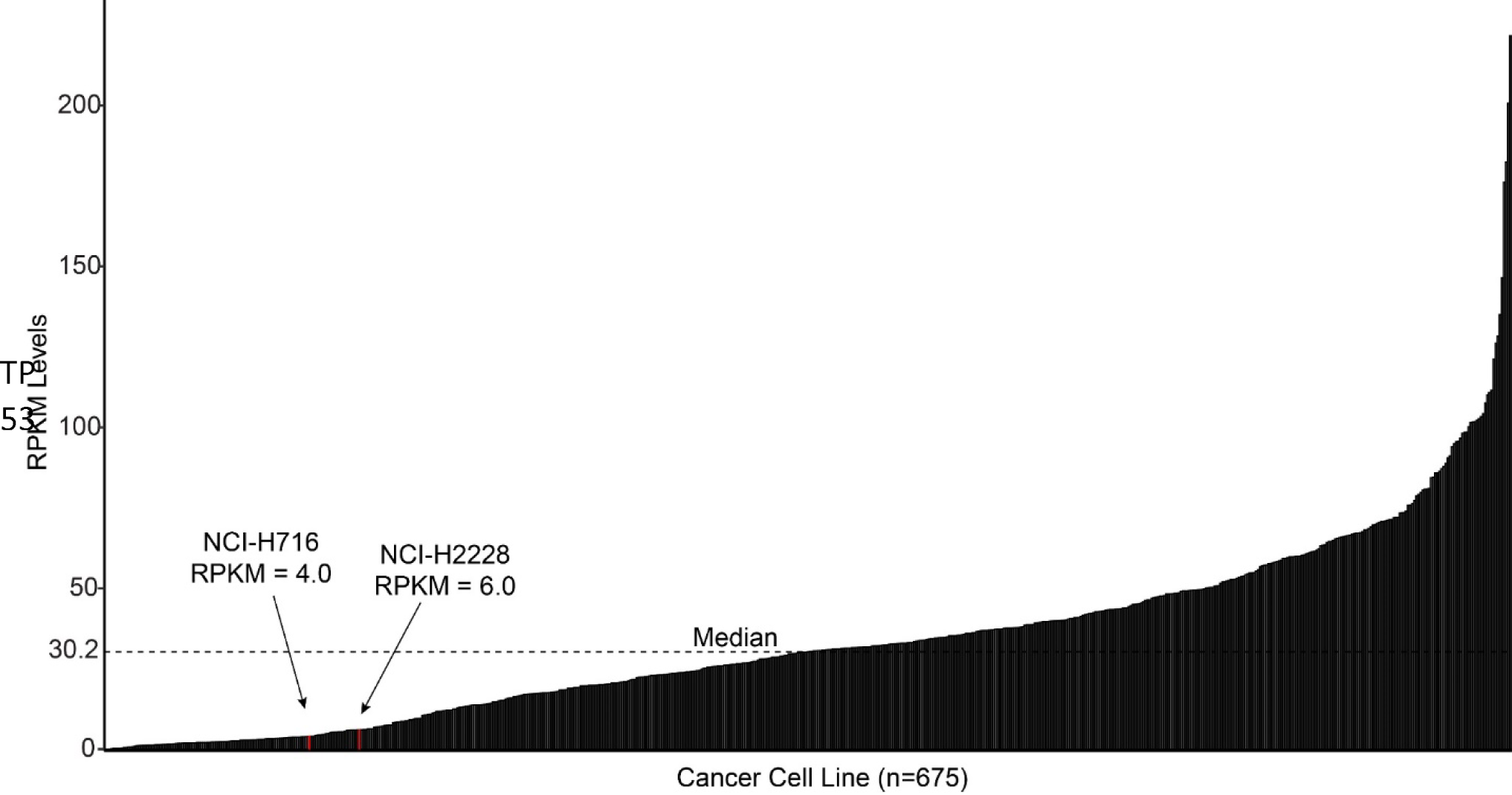
Waterfall plot of publicly available TP53 RNA expression (as shown by FPKM) for 675 cancer cell lines. The two cell lines used in this study are highlighted in red and were in the bottom quartile of *TP53* expression.

**Figure S19:**
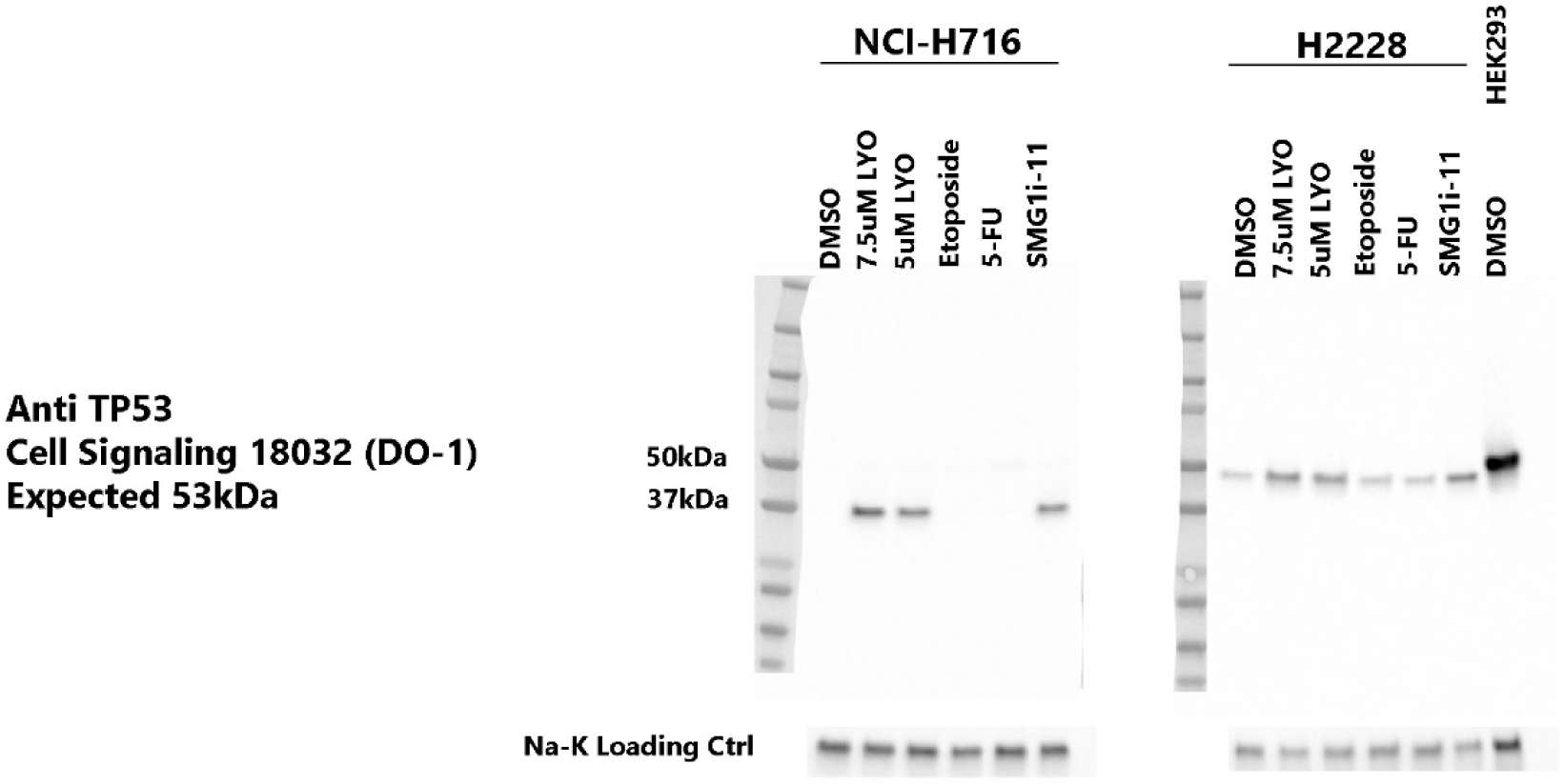
Western blot of TP53 on NCI-H716 and NCI-H2228 cells treated with 5µM or 7.5µM of NMD inhibitor, 1µM SMG1i-11, or 200mg/mL chemotherapy, showing controls related to Figure 4D. LYO is LY3023414 and 5-FU is 5-fluorouracil. HEK293 parent cells are shown as a control (wild type TP53 protein).

**Figure S20:**
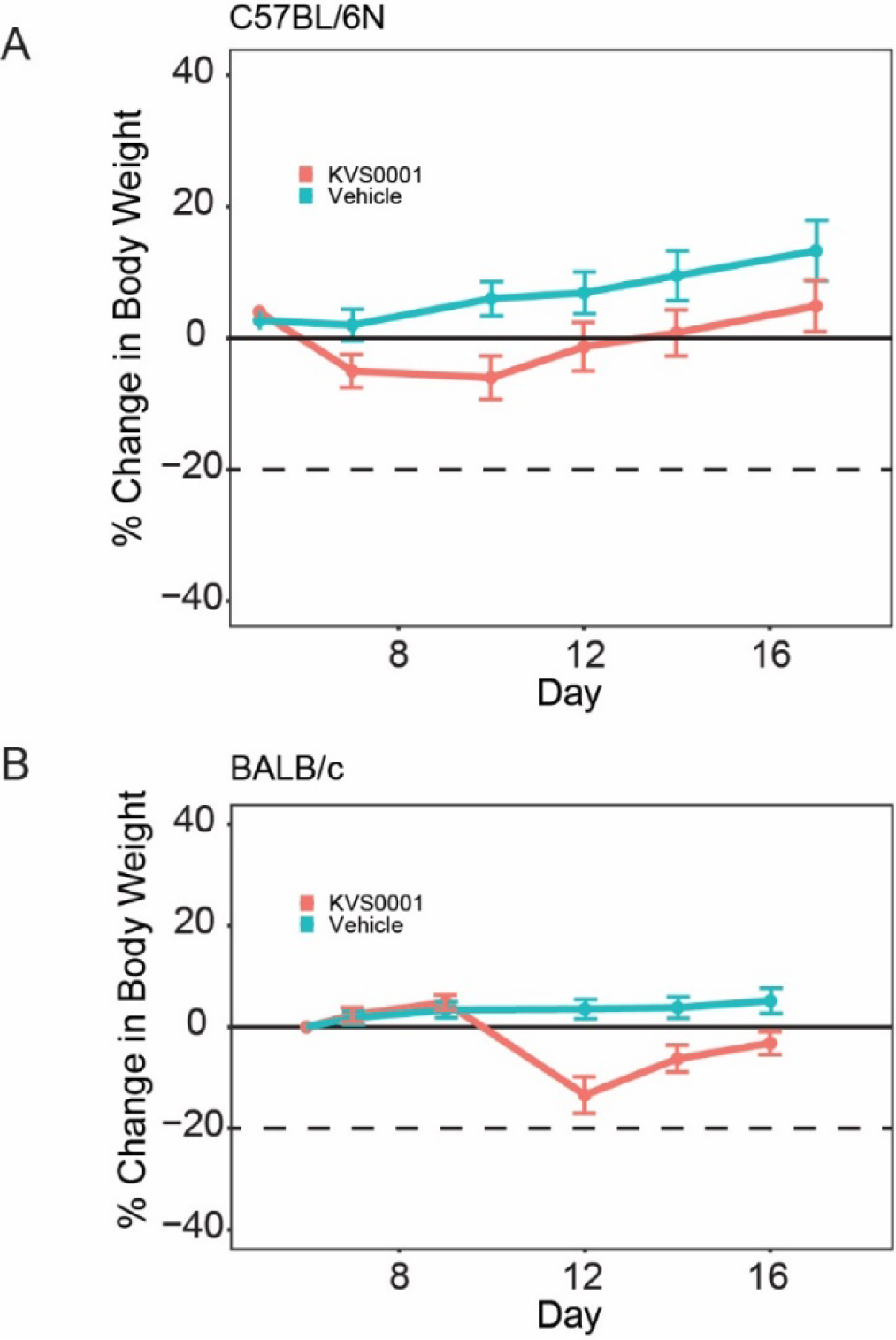
Mouse weights for **A)** C57BL/6N and **B)** BALB/c tumor bearing mice treated with 30mg/kg KVS0001 or vehicle control IP daily. N=8 for all arms.

**Figure S21:**
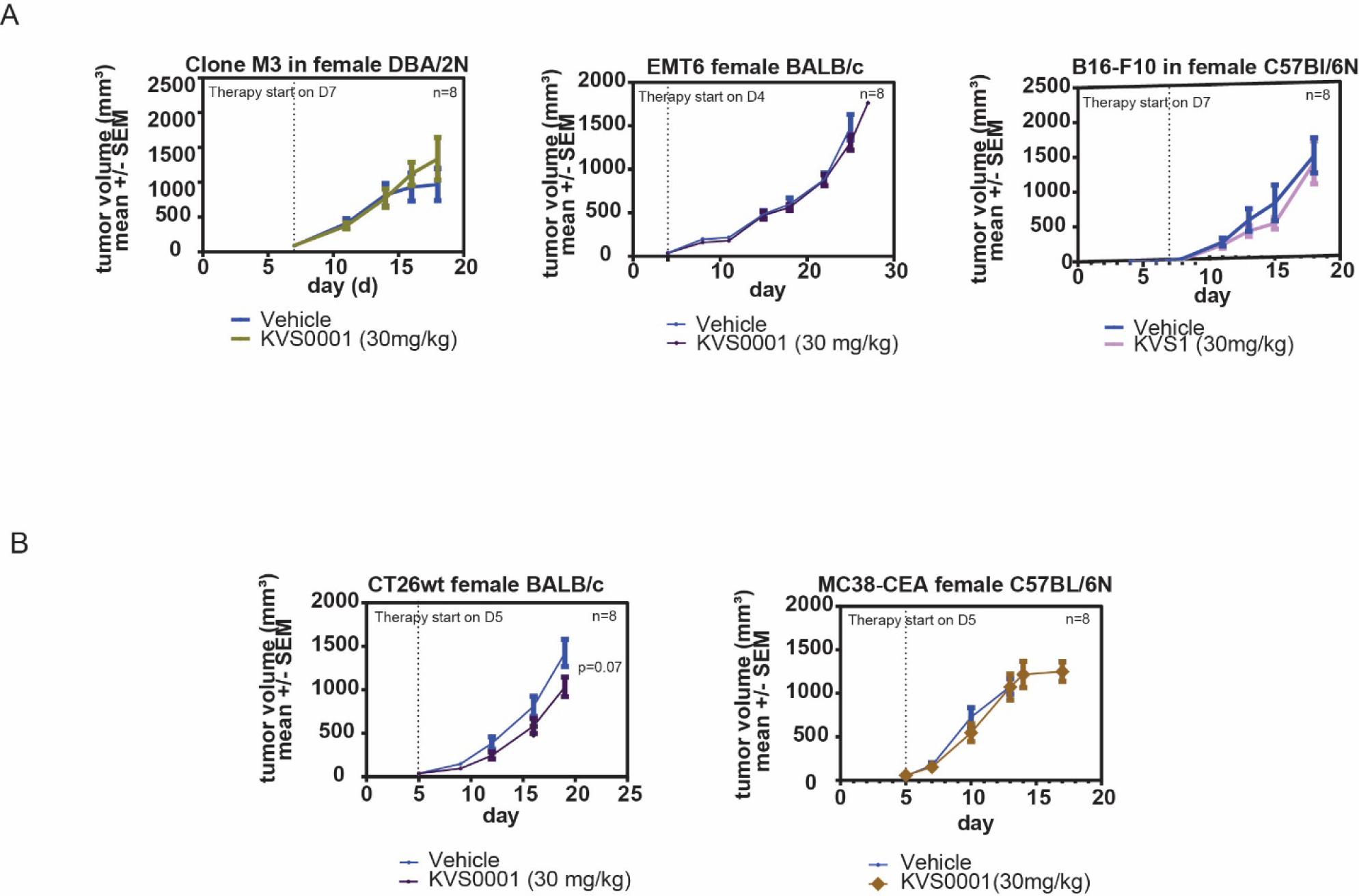
Mouse tumor size as measured by calipers for mammary fat pad placed syngeneic tumor models treated daily with 30mg/kg KVS0001 or vehicle. **A)** Tumors with low/moderate and **B)** high indel mutational loads are shown. No results presented in this supplemental figure are statistically significant. Error bars show 95% confidence intervals.

